# MAP7-driven microtubule remodeling builds the Sertoli apical domain that supports timely meiotic progression

**DOI:** 10.1101/2025.09.16.676497

**Authors:** Koji Kikuchi, Toshihiko Fujimori, Mami Nakagawa, Keisuke Ohta, Ryuki Shimada, Sayoko Fujimura, Kei-ichiro Yasunaga, Shingo Usuki, Naoki Tani, Akira Nakamura, Kimi Araki, Kei-ichiro Ishiguro

## Abstract

How specialized apical membrane domains are assembled *in vivo* remains poorly understood. In the mouse testis, Sertoli cells build a luminal apical domain after birth that organizes the seminiferous epithelium and supports germ cell differentiation, distinct from fetal polarity established during cord formation. Here, we identify the microtubule-associated protein MAP7 as a key regulator of cytoskeletal organization during apical domain formation. MAP7 preferentially localizes to apical microtubules in Sertoli cells, showing limited overlap with stable microtubule markers. Native-tissue imaging indicates that MAP7-decorated microtubules, initially lacking obvious directionality, become increasingly aligned along the tubule axis as the apical domain matures. In *Map7*-deficient testes, microtubule abundance is maintained, but higher-order organization is disrupted and accompanied by persistent luminal F-actin accumulation. These defects compromise apical domain maturation and lumen formation during the first wave of spermatogenesis and are associated with altered tight-junction patterning. Proteomic analysis identifies the non-muscle myosin II heavy chains MYH9 and MYH10 among MAP7-associated proteins. MYH9 becomes enriched at luminal regions where microtubules and F-actin converge, but remains diffuse in *Map7*-deficient Sertoli cells despite ectopic luminal cytoskeletal assemblies, suggesting that MAP7 helps establish a cytoskeletal context that supports spatially restricted NMII enrichment. Single-cell RNA sequencing shows that Sertoli transcriptional differentiation largely proceeds without MAP7, whereas germ cells exhibit delayed meiotic progression with expansion of pachytene-stage subpopulations. Together, these findings establish MAP7-dependent microtubule remodeling as an organizing mechanism for postnatal apical domain maturation and suggest how epithelial cytoskeletal architecture shapes a niche that helps pace developmental progression of neighboring germ cells.

## Introduction

How epithelial cells build and remodel luminal apical architecture *in vivo* remains incompletely understood. In the mouse testis, Sertoli cells provide the epithelial scaffold of seminiferous tubules and support spermatogenesis. During early postnatal development, Sertoli cells undergo a remodeling program that culminates in lumen formation and the emergence of a luminal apical domain. Although Sertoli cells acquire apical–basal polarity during fetal testis cord formation^1^, the luminal-facing apical architecture is built and reorganized after birth. This apical domain shapes tubule organization and supports germ cell differentiation as germ cells move from the basal compartment toward the lumen. In this setting, Sertoli apical structures provide mechanical support and help spatially confine trophic inputs, adhesive interactions, and signaling cues required for maturation^2^. Disruption of apical architecture compromises spermatogenesis and is linked to male infertility^3–6^. However, the cytoskeletal mechanisms that assemble and mature the postnatal luminal apical domain remain poorly defined.

Postnatal maturation of Sertoli apical architecture is accompanied by extensive cytoskeletal reorganization^7^. Actin filaments, major components of ectoplasmic specializations (ES), coordinate adhesion and junctional integrity among Sertoli cells and between Sertoli cells and germ cells^8,9^. Basal ES contributes to the blood–testis barrier (BTB) through actin-based adherens, tight, and gap junctions, whereas apical ES supports Sertoli–spermatid attachment and maturation. Consistently, genetic perturbation of actin regulators in Sertoli cells, including RAC1 and CDC42, disrupts ES organization and compromises postnatal apical architecture^5,6^. In contrast, the role of microtubule remodeling during luminal apical domain assembly has been less clear. Sertoli microtubules form prominent longitudinal bundles along the apical–basal axis and are proposed to support intracellular transport and structural organization. Perturbations of microtubule architecture—such as Sertoli-specific γ-tubulin overexpression^10,11^, dominant-negative EB1 expression^12^, or loss of the microtubule-severing enzyme KATNAL1^13^—lead to spermatogenic defects. These studies implicate microtubules in Sertoli function, but the factors that reorganize apical microtubules as the luminal domain matures *in vivo* remain largely unknown.

Microtubule networks are shaped by microtubule-associated proteins (MAPs)^14,15^. The MAP7 family binds microtubules and recruits kinesin-1, thereby influencing microtubule–motor coupling and network architecture^16–24^. MAP7 proteins can also compete with other MAPs for microtubule occupancy and shape motor distribution in neurons^25,26^. In cultured cells, MAP7 and MAP7D1 guide microtubule plus ends toward the cortex during migration and adhesion^19^. In the mouse testis, MAP7 is expressed in Sertoli and germ cells^27,28^, and *Map7* gene-trap mutants show reduced microtubule bundling in Sertoli cells as well as structural defects in spermatid manchettes^28^. These observations suggest that MAP7 contributes to microtubule organization in the testis, but whether it drives postnatal remodeling of apical microtubules during luminal apical domain formation has remained unresolved.

Here, we define MAP7 function during postnatal Sertoli remodeling and establish how MAP7-dependent microtubule organization supports apical domain maturation. Using *Map7-egfp^KI^* mice, we show that MAP7 associates with a population of apical microtubules in Sertoli cells that shows limited overlap with stable microtubule markers in tissue sections. Native-tissue imaging further indicates that MAP7-decorated bundles, initially lacking obvious directionality, progressively acquire longitudinal orientation as the luminal apical domain matures. In *Map7*-deficient testes, microtubule abundance is largely maintained, but bundle alignment and higher-order organization are disrupted and accompanied by persistent luminal F-actin accumulation. These defects are associated with impaired apical domain maturation, failure of lumen formation, and altered tight-junction patterning. Proteomic analysis identifies the non-muscle myosin II heavy chains MYH9 and MYH10 among MAP7-associated proteins, and MYH9 fails to concentrate at luminal regions where microtubules and F-actin normally converge in *Map7*-deficient Sertoli cells. Finally, scRNA-seq indicates that Sertoli transcriptional differentiation programs are largely preserved without MAP7, whereas germ cells exhibit delayed meiotic progression. Together, these findings link MAP7-dependent microtubule remodeling to postnatal luminal apical domain maturation and to the timing of germ cell development.

## Results

### MAP7 is enriched at the luminal side of Sertoli microtubule bundles

*Map7* mRNA is detected in both Sertoli and germ cells^27,28^, but the protein distribution of MAP7 in the postnatal seminiferous epithelium has not been defined. To visualize endogenous MAP7 *in vivo*, we generated *Map7-egfp* knock-in mice (*Map7-egfp^KI^*; Figures S1A and S1B). Homozygous *Map7-egfp^KI^*mice were viable and fertile (Figure S1C). MAP7-EGFP^KI^ also reproduced the characteristic localization pattern previously reported in the fallopian tube^19^, where it aligned along the planar axis of multiciliated cells on the ovarian (proximal) side (Figure S1D), supporting preservation of MAP7 behavior after EGFP tagging.

We then examined MAP7-EGFP^KI^ in adult testes. Co-staining with the Sertoli cell–enriched β-tubulin isoform TUBB3 revealed that MAP7-EGFP^KI^ decorated Sertoli microtubule bundles and was preferentially enriched toward the luminal side of longitudinal arrays (Figure 1A). These observations indicate that MAP7 exhibits a polarized distribution within Sertoli microtubule networks along the apical–basal axis of the seminiferous epithelium. MAP7-EGFP^KI^ was also detected in elongating spermatids, where it prominently decorated manchette microtubules and overlapped with acetylated tubulin (Figure 1B). Notably, other acetylated microtubules outside the manchette displayed comparatively little MAP7-EGFP^KI^ signal (Figure 1B), suggesting that MAP7 marks a spatially restricted subset of microtubules in elongating spermatids. Antibody staining against MAP7 confirmed the overall localization patterns observed with the knock-in reporter (Figure 1C). Together, these data place MAP7 on two prominent microtubule structures in the seminiferous epithelium—luminally enriched Sertoli bundles and the spermatid manchette—supporting the idea that MAP7 is positioned to influence microtubule architecture during spermatogenesis.

**Figure 1.**
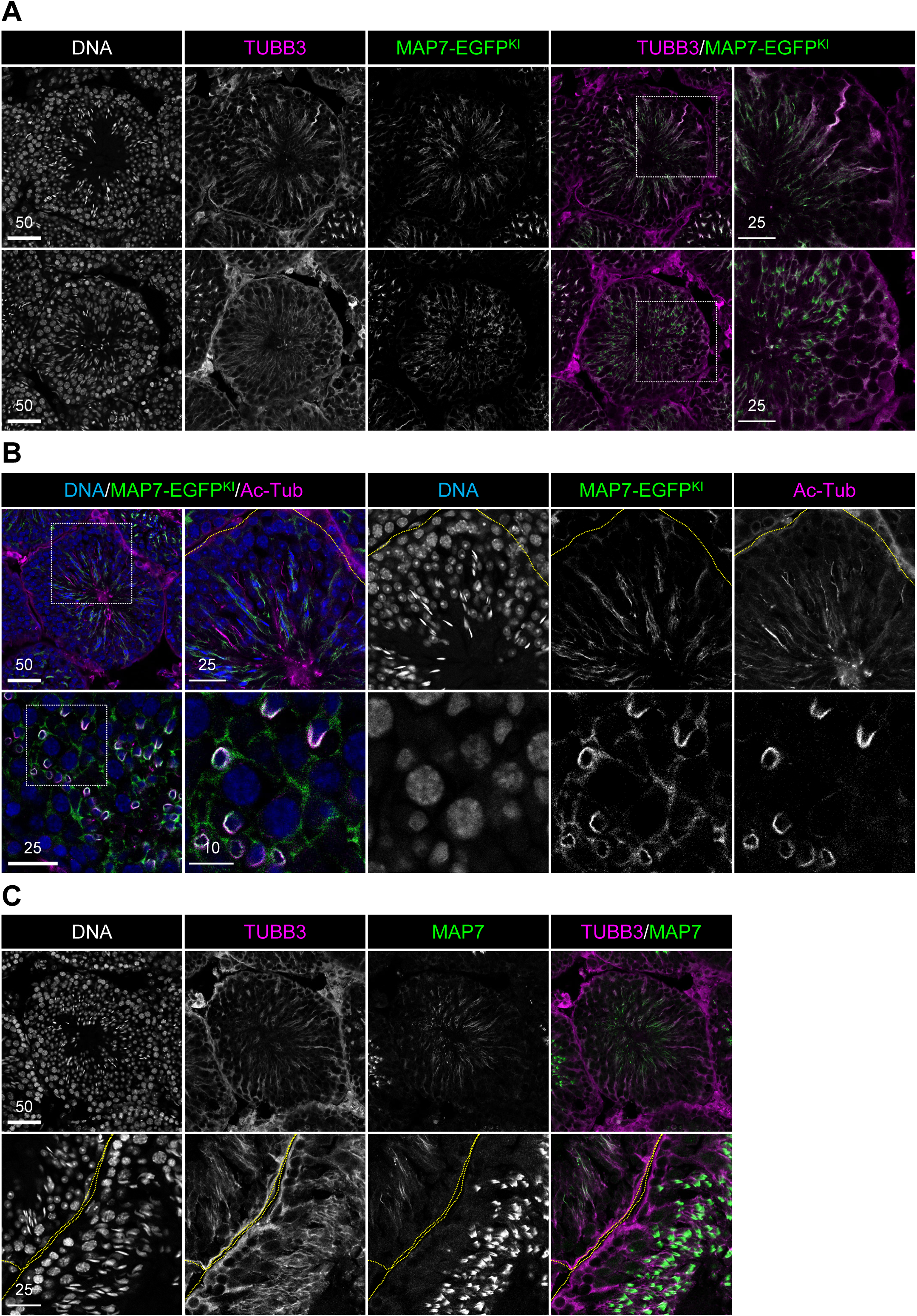
MAP7 is enriched at the luminal side of Sertoli microtubule bundles. **(A)** Seminiferous tubule sections from 8-week-old *Map7-egfp* knock-in (*Map7-egfp^KI^*) mice stained for GFP, βIII-tubulin (TUBB3), and DAPI. TUBB3 marks Sertoli microtubules. Bottom: tubules containing step 9–11 spermatids. Scale bar, 50 or 25 μm. **(B)** Seminiferous tubule sections from 8-week-old *Map7-egfp^KI^* mice stained for GFP, acetylated tubulin, and DAPI. Bottom: higher-magnification view of step 9 spermatids. Scale bar, 50, 25, or 10 μm. **(C)** Seminiferous tubule sections from 8-week-old wild-type mice stained for MAP7, TUBB3, and DAPI. Bottom: tubules containing step 9–11 spermatids. Scale bar, 50 or 25 μm.

### MAP7 is required for postnatal Sertoli remodeling that enables lumen formation

MAP7 is enriched on the luminal side of Sertoli microtubule bundles (Figure 1A), suggesting a role in postnatal epithelial remodeling. Because the previously reported *Map7* gene-trap line was maintained on a mixed background and did not directly confirm complete protein loss^28^, we generated a *Map7* knockout allele on a C57BL/6 background (Figures S2A–C). MAP7 protein was undetectable in *Map7^-/-^*testes by two independent antibodies (Figure S2D). *Map7^-/-^* males showed reduced testis size, spermatogenic defects, and a lack of mature sperm in the cauda epididymis, resulting in complete infertility (Figures S2E and S2F). Heterozygous *Map7^+/-^*littermates, which displayed normal testis morphology, were used as controls.

To define when MAP7 contributes to postnatal tubule remodeling, we examined the first wave of spermatogenesis (P10–P35). Because MAP7 becomes detectable in step 9–11 spermatids around P23, we emphasized earlier stages to reduce germ cell contributions and limit secondary effects associated with later spermatid defects. Although our analysis focused on early postnatal stages, we also examined later ages to assess downstream consequences on tubule architecture. Seminiferous cords were grossly established by P10 in *Map7^-/-^* testes, comparable to *Map7^+/-^* controls (Figure 2A), consistent with MAP7 being dispensable for cord formation. Differences emerged during postnatal remodeling. In *Map7^+/-^* testes, lumina appeared at P15, expanded by P17, and were nearly ubiquitous by P21 (Figures 2A and 2B), consistent with the reported timing of Sertoli maturation and BTB formation^29,30^. In contrast, *Map7^-/-^*tubules largely failed to form lumina between P15 and P28 and often developed abnormal or multilumen structures by P35 (Figures 2A and 2B). These findings indicate that MAP7 is required for normal lumen formation during the first spermatogenic wave.

**Figure 2.**
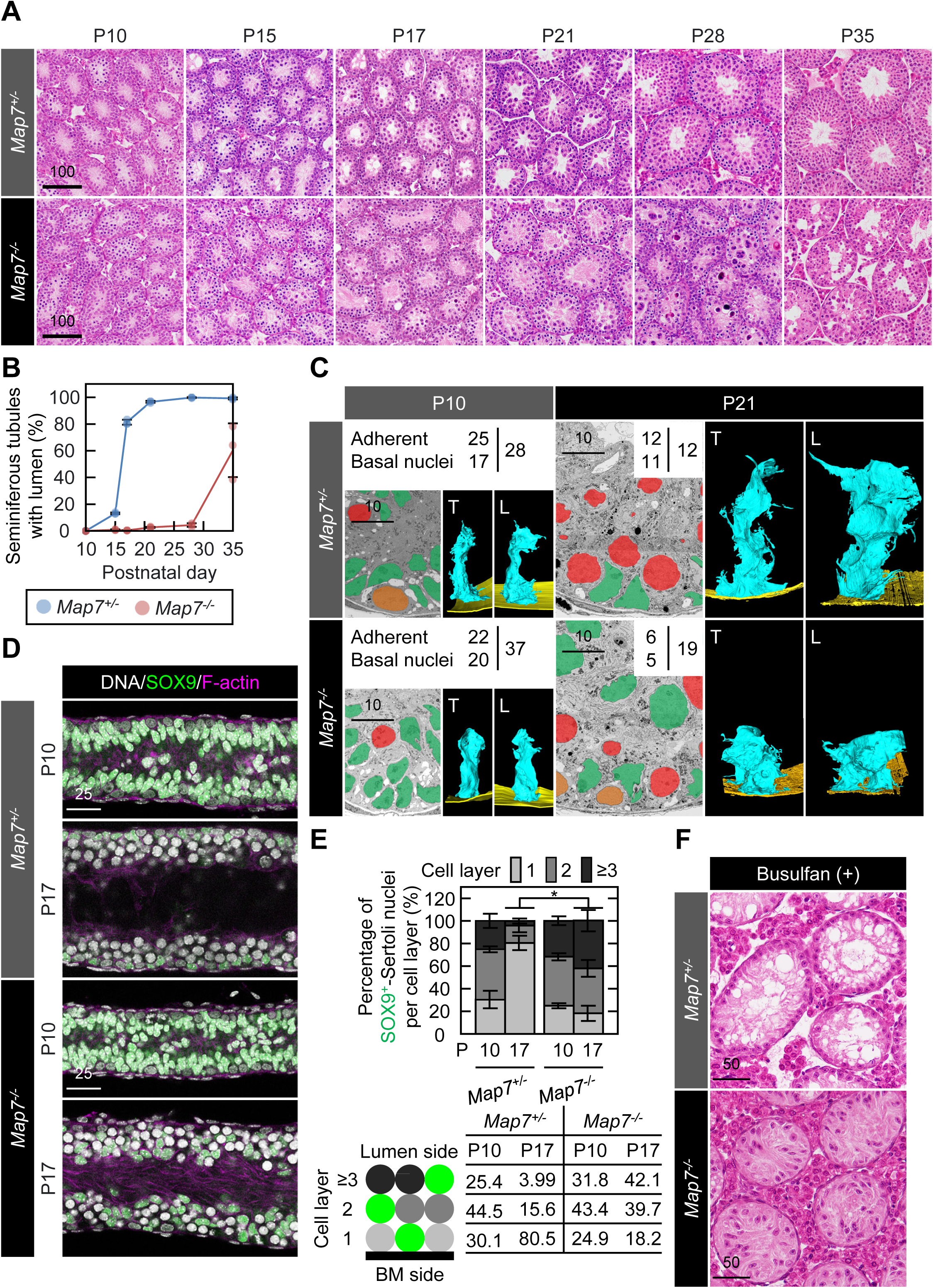
MAP7 is required for postnatal Sertoli remodeling that enables lumen formation. **(A)** Hematoxylin–eosin (HE) staining of testes at postnatal day (P)10, P17, P21, P28, and P35. Three mice per genotype were analyzed. Scale bar, 100 μm. **(B)** Percentage of seminiferous tubules containing a lumen. >140 tubules were quantified across three independent experiments (average ± SD). **(C)** Array tomography (serial-section SEM) of seminiferous tubules at P10 and P21. Serial-section SEM generated 503, 588, 398, and 498 images from *Map7^+/-^*P10, *Map7^-/-^* P10, *Map7^+/-^* P21, and *Map7^-/-^* P21 testes, respectively. Sertoli cells (green), spermatogonia (orange), and spermatocytes (red) were assigned based on nuclear morphology and basement-membrane attachment. Numbers indicate the fraction of Sertoli cells attached to the basement membrane (upper left) or basally positioned (lower left) relative to the total analyzed (right). 3D reconstructions were generated by manual segmentation. Blue and yellow denote Sertoli cells and the basement membrane, respectively; T and L indicate transverse and longitudinal views. Scale bar, 10 μm. **(D)** Basal nuclear positioning and apical actin remodeling in P10 and P17 testes. Tubules were stained with DAPI, SOX9, and phalloidin (F-actin). Scale bar, 25 μm. **(E)** Percentage of SOX9^+^ Sertoli nuclei per layer at P10 and P17. >145 cells per mouse were quantified (≥10 images; three independent experiments; mean ± SD). Two-tailed Student’s *t*-test at P17: *P* < 2.20 × 10^-4^. **(F)** Germ cell–independent maintenance of Sertoli nuclear positioning after busulfan-mediated germ cell depletion. Mice were injected intraperitoneally at P28 (45 μg/g) and analyzed 6 weeks later (N = 4). Scale bar, 50 μm.

Because Sertoli cells interdigitate tightly with germ cells, Sertoli morphology and positioning are difficult to resolve *in situ*. We therefore combined serial sectioning with scanning electron microscopy (SEM) and tracked individual Sertoli cells across consecutive sections to assess basement-membrane contact and nuclear position (Figure 2C). In P10 *Map7^+/-^*testes, most Sertoli cells (89.3%, 25/28) contacted the basement membrane, whereas 60.7% (17/28) of nuclei were basally positioned (Figure 2C). By P21, all Sertoli cells contacted the basement membrane (12/12), and 91.7% (11/12) of nuclei were basally positioned (Figure 2C). SOX9 staining across a larger cohort showed the same developmental shift, with basal nuclear positioning increasing between P10 and P17 (Figures 2D, 2E, and S3A). Over this period, Sertoli cells also extended apically in transverse views and became elongated along the tubule axis (Figure 2C), consistent with coordinated remodeling during luminal domain maturation. These features were disrupted in *Map7^-/-^* testes. At P10, fewer Sertoli cells were attached to the basement membrane (59.5%, 22/37), and 54.1% (20/37) of nuclei were basally positioned (Figure 2C). By P21, fewer Sertoli cells maintained basement-membrane contact (31.6%, 6/19), and only 26.3% (5/19) of nuclei were basally positioned (Figure 2C). SOX9 staining similarly showed persistently low basal positioning, decreasing from 24.9% at P10 to 18.2% at P17 (Figures 2D, 2E, and S3A), and the defect remained evident in adulthood (Figures S3B and S3C). Even among Sertoli cells that retained basement-membrane contact at P21, apical extension and longitudinal elongation appeared blunted (Figure 2C). Actin remodeling was also altered: in controls, luminal F-actin was detectable at P10 but diminished by P17 as lumina formed (Figures 2D and S3A), whereas luminal F-actin persisted at P17 in *Map7^-/-^* testes (Figures 2D and S3A). Thus, MAP7 loss disrupts postnatal Sertoli remodeling, including basement-membrane attachment, basal nuclear positioning, and coordinated changes in cell shape and actin-based luminal organization.

Because Sertoli organization could in principle be influenced by germ cells^31–35^, we asked whether germ cell input is required to maintain basal positioning of Sertoli nuclei once established. We depleted spermatogonial stem cells (SSCs) with busulfan at P28 and analyzed testes 6 weeks later, when germ cell lineages were effectively eliminated. After treatment, *Map7^+/-^* mice showed testis-to-body weight ratios comparable to untreated *Map7^-/-^* mice (Figure S3D), and hematoxylin–eosin (HE) staining confirmed germ cell loss (Figures 2F and S3E). In *Map7^+/-^*testes, Sertoli nuclei remained predominantly basally positioned despite germ cell depletion. In contrast, Sertoli nuclei in *Map7^-/-^* testes frequently shifted toward the lumen under the same conditions (Figures 2F and S3E). These results indicate that the nuclear positioning defect in *Map7*-deficient Sertoli cells is not simply secondary to the presence of ongoing germ cell lineages.

Collectively, these results show that MAP7 is required for postnatal Sertoli remodeling that establishes apical–basal organization and supports lumen formation. In *Map7^-/-^* testes, reduced basement-membrane attachment, altered nuclear positioning, and impaired cell elongation are accompanied by persistent luminal F-actin accumulation and defective lumen expansion, resulting in sustained disruption of seminiferous tubule architecture.

### MAP7 promotes microtubule alignment during postnatal apical domain maturation

To link these remodeling defects to cytoskeletal reorganization, we next examined how microtubules are remodeled during postnatal apical domain maturation and lumen formation. Although Sertoli cells acquire apical–basal features during fetal testis cord formation, the luminal apical architecture characteristic of the seminiferous epithelium is assembled after birth. We therefore focused on the P10–P17 transition, when apical maturation and lumen formation proceed, and examined microtubules and F-actin at P10 and P17 to capture cytoskeletal reorganization during this window. At P10, TUBB3-positive microtubules and F-actin were enriched on the luminal side of seminiferous tubules, and MAP7-EGFP^KI^ signal accumulated in the same inner region (Figure 3A, top). By P17, the prominent luminal enrichment was largely resolved: microtubules became organized within elongating Sertoli processes along the luminal–basal axis, whereas F-actin concentrated at the basal ES adjacent to the basement membrane (Figure 3A, bottom). At this stage, MAP7 showed a polarized distribution along Sertoli microtubules, with enrichment toward the luminal side, consistent with the pattern observed in adult testes (Figures 1A, 1C, and 3A, bottom). To assess the orientation of MAP7-decorated microtubules in native tissue, we imaged freshly dissected seminiferous tubules from *Map7-egfp^KI^* mice. At P10, apical MAP7-EGFP^KI^–positive bundles did not exhibit a consistent orientation and appeared variably oriented (Figure 3B, left). By P17, MAP7-decorated apical microtubules appeared predominantly longitudinally aligned along the tubule axis (Figure 3B, right). This change parallels the elongation and longitudinal orientation of apical Sertoli processes observed by SEM between P10 and P21 (Figure 2C). These observations suggest that postnatal apical domain maturation is accompanied by progressive ordering of MAP7-marked apical microtubule bundles.

**Figure 3.**
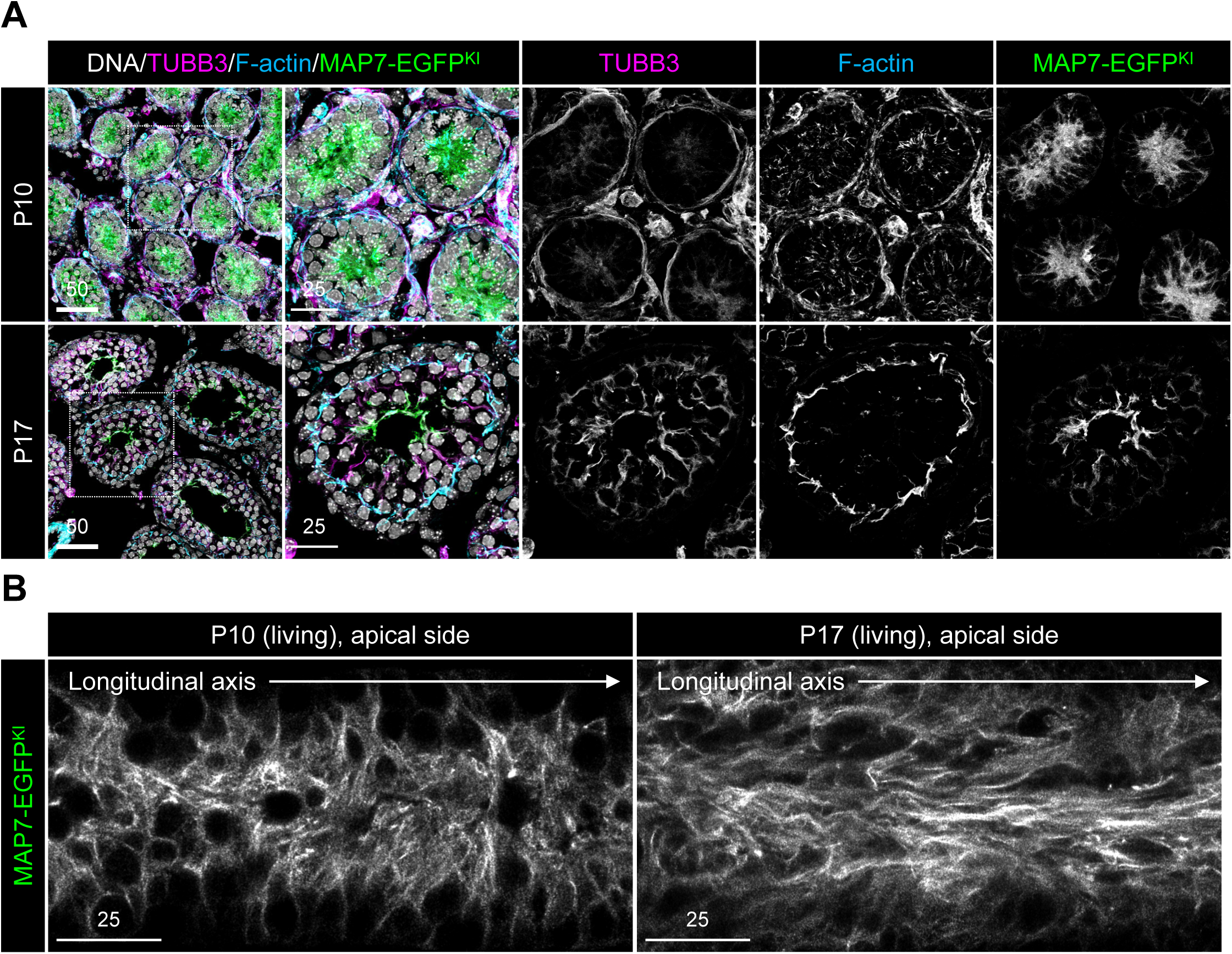
Cytoskeletal remodeling and MAP7 localization during postnatal apical domain maturation. **(A)** Seminiferous tubule sections from *Map7-egfp^KI^* mice at P10 and P17 stained for GFP, βIII-tubulin (TUBB3), phalloidin (F-actin), and DAPI. Scale bar, 50 μm. **(B)** Confocal imaging of freshly dissected seminiferous tubules from *Map7-egfp^KI^* mice at P10 (left) and P17 (right). Scale bar, 25 μm.

We next examined how *Map7* deletion affects cytoskeletal remodeling during this postnatal window. In *Map7^+/-^* testes at P17, the luminal microtubule enrichment prominent at P10 was no longer apparent, and F-actin was largely confined to the basal ES (Figure 4A, top). In *Map7^-/-^* testes, however, luminal microtubule signals persisted at P17, resembling the P10 configuration. Microtubule staining intensity was broadly preserved, but bundle alignment along the luminal–basal axis was disrupted (Figure 4A, bottom). Luminal F-actin accumulation likewise persisted (Figure 4A, bottom). Across tubules, the frequency of ectopic inner-side accumulation of TUBB3 and F-actin was similar to the fraction of tubules that failed to form a lumen (Figures 2A, 2B, and 4A), consistent with an association between persistence of the early cytoskeletal configuration and failed lumen formation. Similar features were observed in adult testes. In controls, Sertoli microtubule bundles extended along the luminal–basal axis and apical actin structures were organized normally (Figure S4A, top). In *Map7^-/-^*testes, microtubule abundance remained comparable, but alignment defects persisted and apical actin organization was replaced by ectopic structures (Figure S4A, bottom). These observations are consistent with a model in which MAP7 promotes higher-order organization and alignment of apical microtubule bundles without primarily reducing overall microtubule abundance.

**Figure 4.**
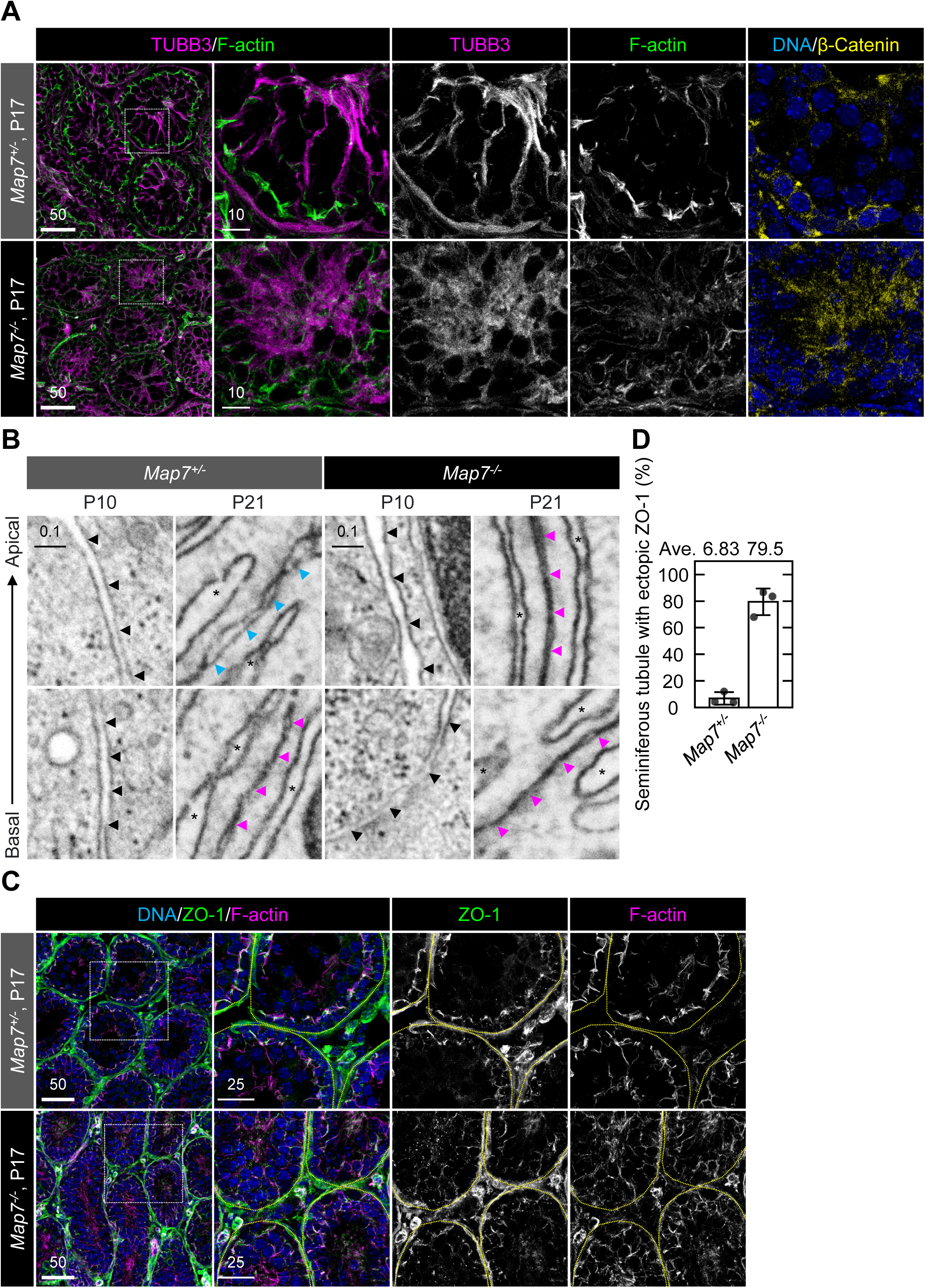
MAP7 promotes microtubule alignment during postnatal apical domain maturation. **(A)** Microtubule bundle organization in Sertoli cells at P17. Three-dimensional projection images of seminiferous tubule sections stained for βIII-tubulin (TUBB3), phalloidin (F-actin), β-catenin, and DAPI. The fraction of *Map7^-/-^* tubules showing ectopic inner-side accumulation of TUBB3/F-actin/β-catenin matched the fraction lacking lumen formation (Figures 2A and 2B). Scale bars, 50 or 25 μm. **(B)** Ultrastructural analysis of Sertoli–Sertoli junctions by high-resolution SEM at P10 and P21. The black arrow indicates the Sertoli–Sertoli boundary at P10; magenta and blue arrows indicate tight junctions and adherens junction–like structures at P21, respectively. Asterisks mark the endoplasmic reticulum. Scale bar, 0.1 μm. **(C)** Tight junction positioning at P17 visualized by three-dimensional projections of seminiferous tubules stained for ZO-1, phalloidin (F-actin), and DAPI. Scale bars, 50 or 25 μm. **(D)** Quantification of tubules with ectopic ZO-1 localization based on images in (C). Data are average ± SD from >45 tubules across three independent experiments. Student’s *t*-test, *P* < 0.002.

Given the MAP7-dependent changes in apical cytoskeletal organization, we asked whether junctional architecture is also affected during postnatal maturation. In the testis, tight junctions form at the basal side of Sertoli cells and contribute to BTB assembly, thereby partitioning the seminiferous epithelium into basal and luminal compartments. We therefore examined Sertoli–Sertoli junctions by SEM. In P21 *Map7^+/-^* testes, electron-dense regions along the basal side of Sertoli–Sertoli contacts were consistent with tight junctions closely associated with the endoplasmic reticulum, as reported previously^36^ (Figures 4B and S4B). Immediately apical to these sites, we observed junctional regions that were also ER-associated but displayed wider intercellular spacing than tight junctions. This morphology is consistent with adherens junction–like structures, reflecting looser membrane apposition; however, these regions could not be definitively distinguished from gap junctions or desmosomes based on ultrastructure alone (Figures 4B and S4B). At P10, both *Map7^+/-^* and *Map7^-/-^* testes lacked clearly defined tight junctions and adherens junction–like structures (Figures 4B and S4B). By P21, *Map7^-/-^*Sertoli cells displayed expanded basal tight junction regions but failed to establish adherens junction–like structures (Figures 4B and S4B). Consistent with this ultrastructural phenotype, the tight junction marker ZO-1 localized predominantly to the basal ES, marked by F-actin, in *Map7^+/-^*testes, whereas in *Map7^-/-^* testes ZO-1 mislocalized to ectopic F-actin–rich regions (Figures 4C and 4D). β-catenin, a core component of adherens junctions, colocalized with F-actin near the basal ES in *Map7^+/-^* testes, but accumulated toward the luminal side in *Map7^-/-^*testes (Figures 4A and S4A). Together, these results indicate that loss of MAP7 is associated with altered spatial organization of junctional assemblies during postnatal Sertoli maturation.

### MAP7 coordinates microtubules with actomyosin-associated regulators during postnatal apical domain maturation

MAP7-dependent microtubule remodeling accompanies postnatal apical domain maturation and is coupled to changes in junctional patterning, raising the possibility that MAP7 links microtubules to other cytoskeletal systems. To identify candidate effectors, we performed immunoprecipitation–mass spectrometry (IP–MS) to define MAP7-associated proteins. We analyzed two datasets: *MAP7-egfp^KI^* HeLa cells (Figure 5A) and testes from P17–P20 *Map7-egfp^KI^* mice (Figure 5B). The HeLa dataset served as a reference for robust MAP7 interactions, whereas the testis dataset captured partners present during postnatal Sertoli remodeling. The kinesin-1 heavy chain KIF5B, a well-established MAP7 interactor, was detected in both datasets (Figures 5A–C), supporting purification specificity. Comparative analysis identified 22 proteins selectively enriched in MAP7 immunoprecipitates from P17–P20 testes (Figure 5C). Functional enrichment analysis (Metascape) highlighted categories including motor proteins and Rho GTPase effectors (Figure 5D). Among these candidates, we focused on cytoskeletal regulators (Figure 5E), including the Sertoli β-tubulin isoform TUBB3, non-muscle myosin II (NMII) heavy chains MYH9/MYH10, and the intermediate filament protein vimentin (VIM), consistent with MAP7 engaging multiple cytoskeletal modules during apical domain maturation.

**Figure 5.**
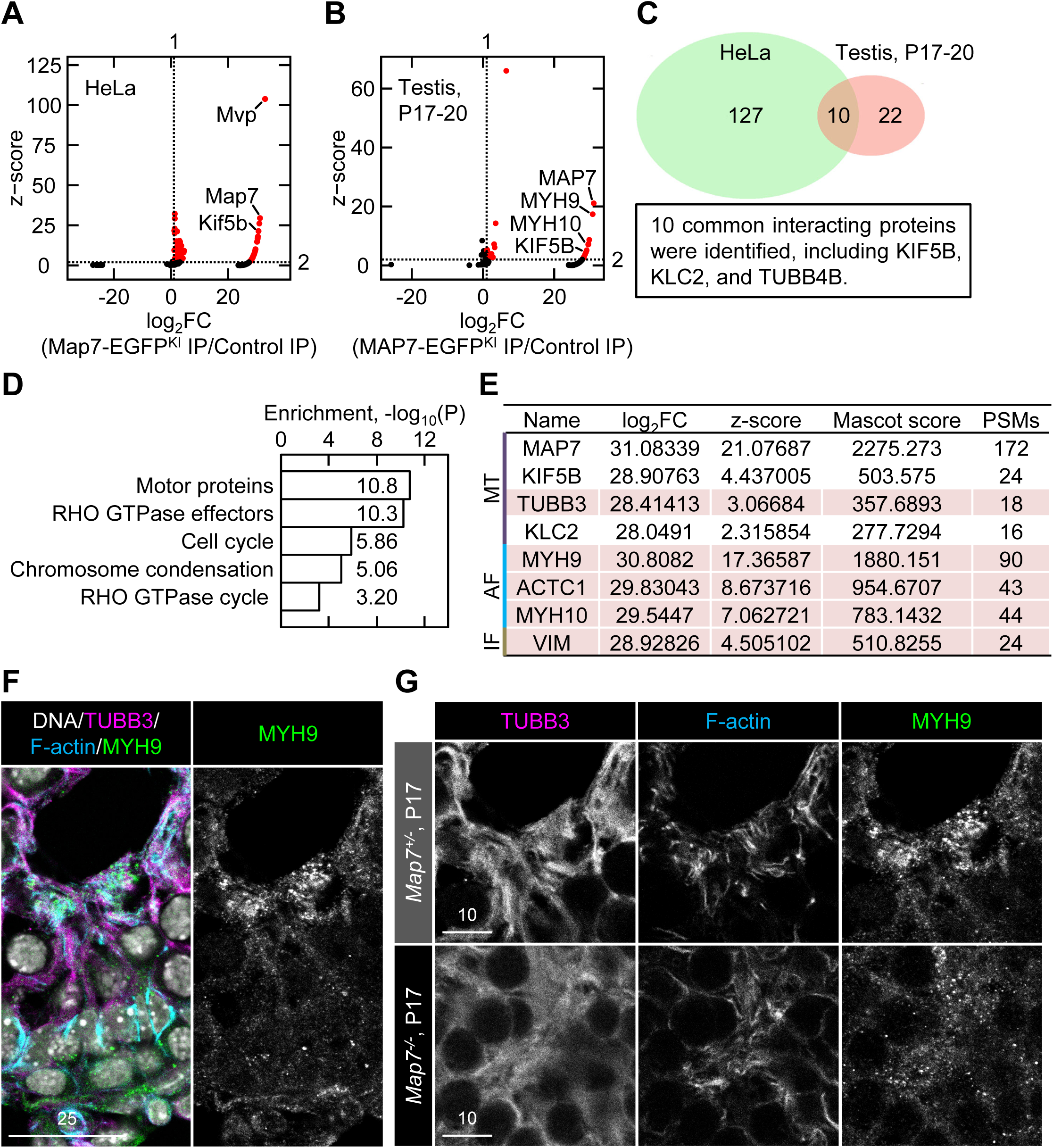
MAP7 coordinates microtubules with actomyosin-associated regulators during postnatal apical domain maturation. **(A)** IP–MS analysis of MAP7 interactors in *MAP7-egfp^KI^* HeLa cells. Wild-type lysates served as negative controls. Volcano plot shows proteins identified after excluding keratin-and immunoglobulin-derived species. Enrichment was quantified using Mascot protein scores; proteins with log2 fold change > 1 and z-score > 2 are highlighted in red. High-scoring proteins, including MVP, MAP7, and KIF5B, are labeled. **(B)** IP–MS analysis of *Map7-egfp^KI^*testes at P17–P20, with age-matched wild-type lysates as controls. Proteins meeting the same criteria (log2 fold change > 1; z-score > 2) are highlighted in red, with MAP7, MYH9, MYH10, and KIF5B labeled. **(C)** Venn diagram comparing proteins co-purified in HeLa cells and P17–P20 testes. Ten proteins were shared between datasets, including KIF5B, KLC2, and the β-tubulin isoform TUBB4B. **(D)** Metascape enrichment analysis of 22 proteins uniquely detected in MAP7 immunoprecipitates from testes and annotated as Sertoli-enriched. **(E)** Cytoskeleton-associated proteins absent from control IPs are shown, with Sertoli-enriched proteins highlighted in red. Proteins are categorized by association with microtubules (MT), actin filaments (AF), or intermediate filaments (IF). Mascot scores indicate identification confidence, and peptide-spectrum matches (PSMs) indicate the number of confidently matched spectra. **(F)** Immunofluorescence of testes from P17 *Map7^+/-^* mice stained for TUBB3, MYH9, phalloidin (F-actin), and DAPI. Scale bar, 25 μm. **(G)** Higher-magnification views of MYH9 localization in *Map7^+/-^*and *Map7^-/-^* Sertoli cells. Grayscale channels for TUBB3, MYH9, and F-actin are shown; corresponding lower-magnification images are shown in Figure S5. Scale bar, 10 μm.

We prioritized MYH9 because of its strong enrichment and known role in actomyosin-driven remodeling. In control testes, MYH9 showed a clear spatial bias: it was enriched at luminal regions where microtubules and F-actin converged, but was low in regions lacking either component (Figures 5F and S5). In *Map7^-/-^* testes, microtubule and F-actin assemblies persisted in luminal regions where they are normally cleared during maturation; however, MYH9 did not accumulate at these ectopic structures and instead remained diffuse throughout the cytoplasm (Figures 5G and S5). Together, these observations support that MAP7 is required for normal luminal enrichment of MYH9 and suggest that MAP7-dependent microtubule organization provides a spatial context that facilitates NMII positioning during postnatal apical domain maturation.

### Sertoli cell transcriptional states are largely preserved without MAP7

Our data indicate that MAP7 is required for postnatal cytoskeletal remodeling and apical domain maturation in Sertoli cells, raising the question of whether MAP7 also influences Sertoli functional differentiation at the transcriptional level. To address this, we performed scRNA-seq of Sertoli cells at P19. GFP-positive Sertoli cells were isolated by FACS using a *Sox9-ires-gfp* allele (Figure S6A) and profiled with the 10x Genomics platform. Sertoli cells from wild-type and *Map7^+/-^* littermates were transcriptionally indistinguishable and were therefore combined as a single control group (Figure S6B). Although the GFP-positive sorted fraction contained a subset of cells with germ cell–like transcriptomes, we stringently defined Sertoli cells *in silico* by selecting cells expressing canonical Sertoli markers (*Gata4*, *Sox9*, *Kitl*, and *Dhh*) and excluding cells expressing germ cell markers (e.g., *Ddx4*). This yielded 311 control and 434 *Map7^-/-^* Sertoli cells for downstream analyses (Figure S6B).

UMAP clustering resolved six transcriptionally distinct Sertoli cell states (C0–C5) in both genotypes (Figure 6A). Gene set enrichment analysis (Metascape) suggested that these clusters are consistent with conserved specialization programs (Figures 6B and S7). C0 showed a stress-response signature. C1 was metabolically active, with enrichment for fructose metabolism and antioxidant capacity. C2 was enriched for signaling-responsive and apical/junction-associated gene signatures. C3 exhibited a cytoskeletal/TGFβ-associated program consistent with ECM remodeling and proliferation control. C4 was enriched for ECM-related genes consistent with structural specialization, whereas C5 showed a high-energy, stress-adapted program that co-upregulated carbon metabolism and stress-response pathways. These programs were present in both genotypes, and no aberrant Sertoli state emerged in *Map7^-/-^*testes (Figures 6A and 6E).

**Figure 6.**
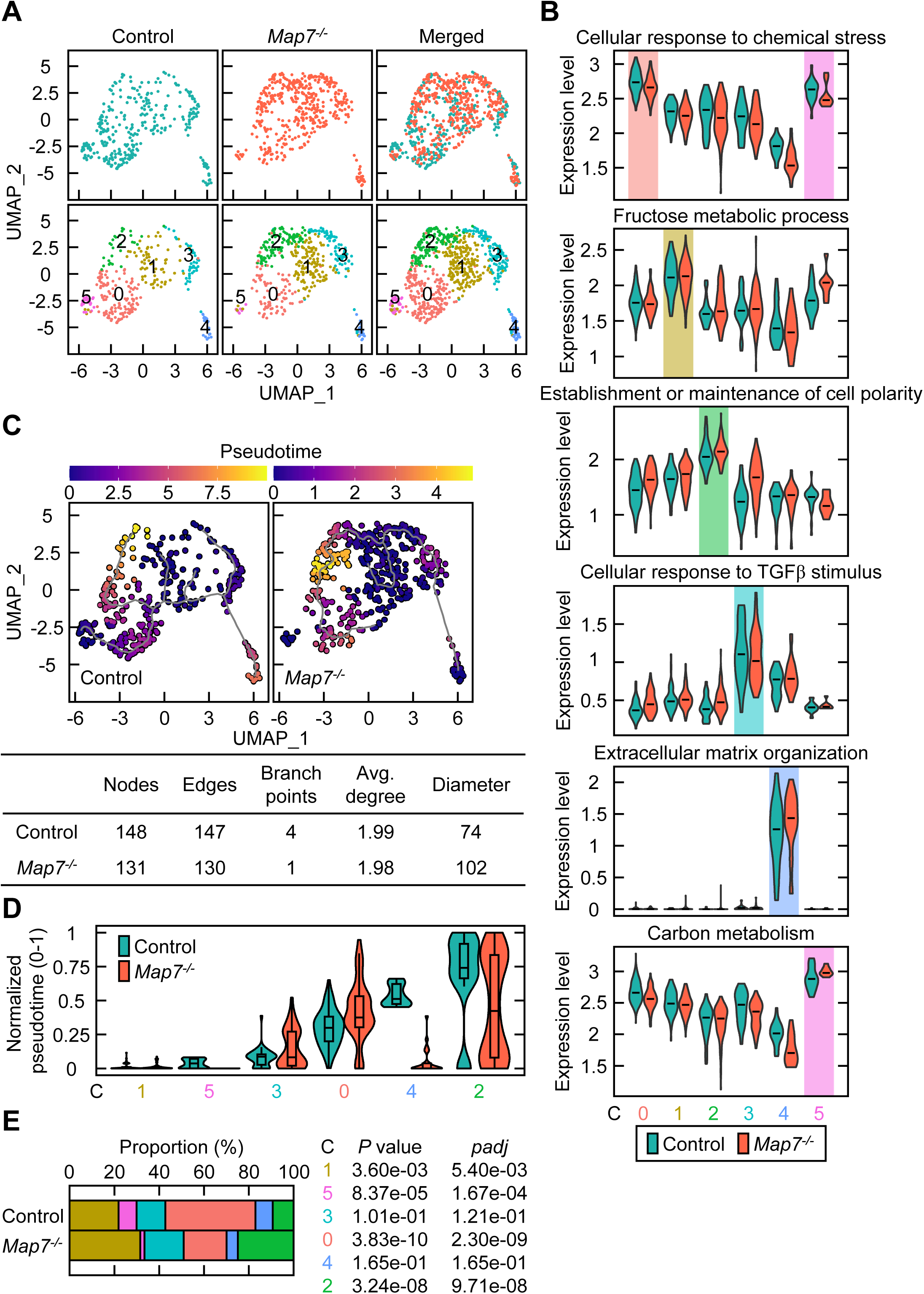
Sertoli cell transcriptional states are largely preserved without MAP7. **(A)** UMAP visualization of transcriptomic profiles from Sertoli cells isolated from P19 testes of the indicated genotypes. The control sample was generated by pooling testes from one wild-type and two *Map7^+/-^* littermates; *Map7^-/-^* littermates were analyzed in parallel. Clusters were defined based on transcriptional similarity. **(B)** Violin plots showing representative cluster-enriched gene sets derived from the top 100 differentially expressed genes per cluster, identified by Metascape enrichment analysis. **(C)** UMAPs for control and *Map7^-/-^* Sertoli cells plotted separately (Seurat). Using the same datasets, pseudotime trajectories were inferred with Monocle3. Root selection was based on shared_score, specificity_score, and entropy_score, resulting in cluster 1 being selected as the root. Graph statistics (nodes, edges, branch points, average degree, and diameter) are shown below the corresponding UMAPs. **(D)** Distribution of normalized pseudotime values (0–1) across clusters for control and *Map7^-/-^*Sertoli cells. Violin plots with embedded boxplots highlight shifts in pseudotime positioning between genotypes. **(E)** Proportions of Sertoli cells assigned to each cluster in control and *Map7^-/-^* testes. Statistical differences were assessed using Fisher’s exact test; *P* values and adjusted *P* values (*padj*) are shown.

To assess relationships among these states, we reconstructed transcriptional ordering using Monocle3. In controls, the inferred ordering formed a continuous maturation trajectory across clusters, proceeding from C1 → C5 → C3 → C0 → C4 → C2 (Figure 6D). *Map7^-/^*^-^Sertoli cells occupied the same trajectory, with modest shifts in the relative placement of intermediate states, whereas the terminal state (C2) remained shared between genotypes (Figure 6D). Graph topology was similarly preserved: Monocle3 principal graphs showed nearly identical average node degrees between genotypes, and branch points and graph diameter differed only minimally (Figure 6C). Consistent with this, cluster proportions and overall distribution across UMAP space were broadly similar (Figures 6A and 6E). These results indicate that MAP7 is not required to establish the overall transcriptional landscape of postnatal Sertoli cell states, but they reveal modest genotype-associated shifts in positioning along pseudotime.

### Sertoli apical domain supports timely pachytene progression during meiosis

Because *Map7* loss disrupts postnatal apical domain formation and junctional patterning, we asked how these architectural defects affect the Sertoli cell microenvironment that supports germ cell development. We first examined whether the SSC pool is maintained in *Map7^-/-^* testes. Given that Sertoli adhesion contributes to SSC niche function^37,38^, we quantified GFRα1^+^ SSCs in 8-week-old *Map7^+/-^* and *Map7^-/-^* mice. GFRα1^+^ SSC numbers were comparable between genotypes (Figures 7A and 7B), indicating that SSC maintenance is largely preserved despite defects in apical domain formation and junctional organization. We next assessed spermatogenic progression in adult testes using ZFP541, which marks pachytene spermatocytes through round spermatids^39^. Cell populations were quantified based on nuclear morphology and SYCP3 co-expression. In *Map7^-/-^*testes, both pachytene spermatocytes and round spermatids were reduced, with a more pronounced decrease in round spermatids (Figures 7C, 7D, and S8A). TUNEL staining did not reveal a detectable increase in apoptosis in either genotype (Figure S8B), suggesting that reduced round spermatids are unlikely to be explained by overt apoptosis and are consistent with delayed progression into the round spermatid stage.

**Figure 7.**
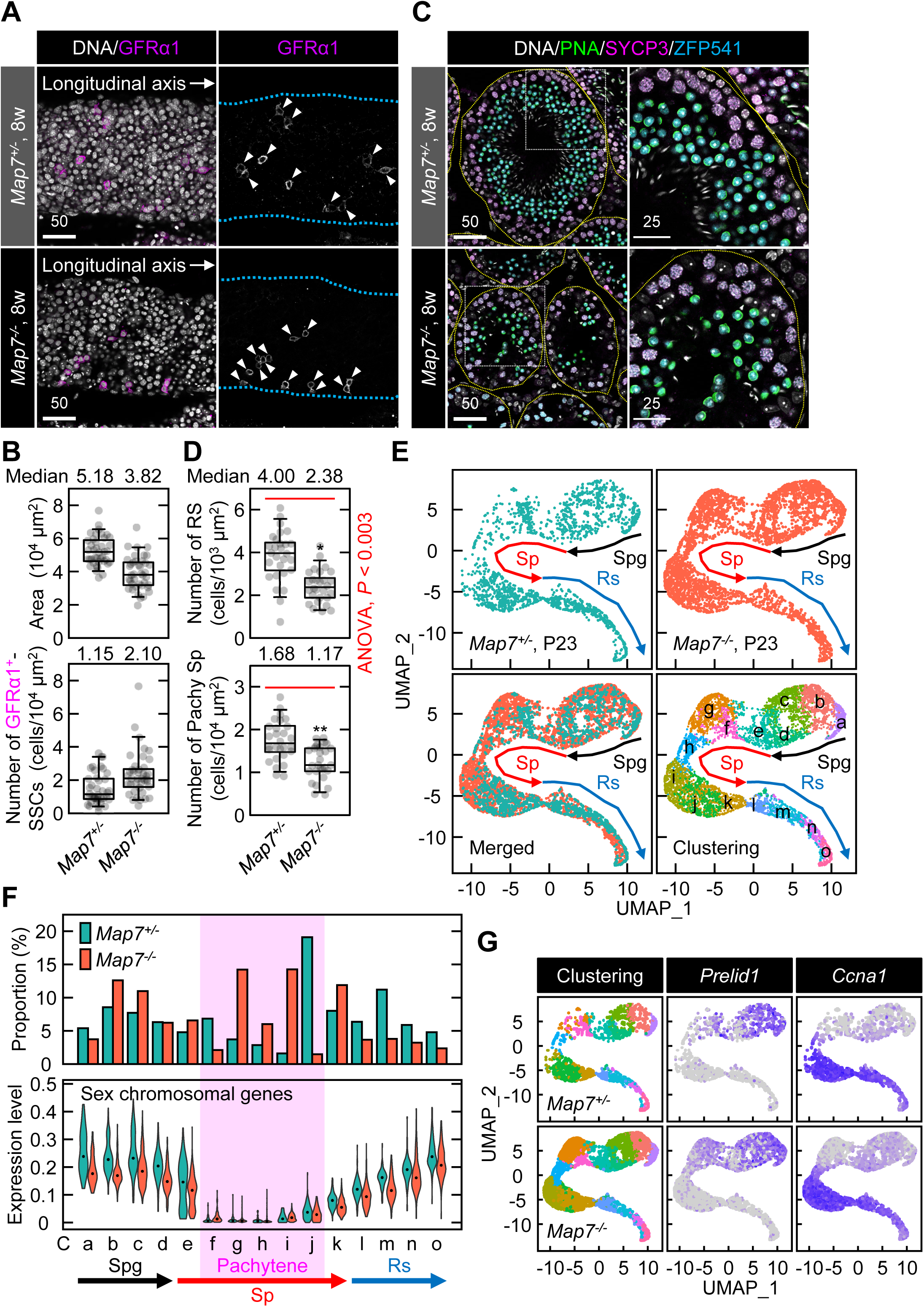
Sertoli apical domain supports timely pachytene progression during meiosis. **(B)** Whole-mount immunofluorescence of seminiferous tubules from 8-week-old mice showing GFRα1^+^ spermatogonial stem cells (SSCs). Tubules were stained for GFRα1 and DAPI and imaged basally by z-sectioning. Scale bar, 50 μm. **(C)** Quantification of GFRα1^+^ SSCs per 10^4^ μm^2^ based on basal tubule area. Data were collected from 40 tubules per genotype across three independent experiments. **(D)** Immunofluorescence of ZFP541^+^ round spermatids and SYCP3^+^/ZFP541^+^ pachytene spermatocytes in testis sections from 8-week-old mice. Sections were stained with PNA, SYCP3, ZFP541, and DAPI. Scale bars, 50 and 25 μm. **(E)** Quantification of ZFP541^+^ round spermatids (RS) and SYCP3^+^/ZFP541^+^ pachytene spermatocytes (Pachy Sp) per 10^3^ μm^2^ from 28 images per genotype across three independent experiments. One-way ANOVA (*P* < 0.003) followed by Student’s *t*-test: *, *P* < 4 × 10^-6^; **, *P* < 4 × 10^-5^. **(F)** UMAP visualization of scRNA-seq profiles from germ cells isolated from P23 testes. Clusters were defined based on gene expression, and arrows indicate inferred differentiation trajectories. Spg, Sp, and RS denote spermatogonia, spermatocytes, and round spermatids. **(G)** Proportions of germ cell populations across clusters. Spermatocyte sub-stages were further resolved by sex-chromosomal gene silencing, shown as violin plots. **(H)** UMAPs of the indicated genotypes (left) and expression of *Prelid1* (early pachytene) and *Ccna1* (late pachytene) (middle and right), supporting classification of pachytene sub-states and highlighting altered progression in *Map7^-/-^* testes.

To profile germ cell consequences at higher resolution, we performed scRNA-seq at P23, before the appearance of step 9–11 spermatids (Figure S8C). This timing enabled focused analysis of earlier spermatogenic populations. At P23, phenotypes observed in *Map7^-/-^*testes—including luminal closure, abnormal microtubule organization, and ectopic accumulation of F-actin and β-catenin—remained evident (Figures S8C and S8D). Whole testes from *Map7^+/-^* and *Map7^-/-^* mice were dissociated and subjected to scRNA-seq using the 10x Genomics platform. Germ cells were identified by *Ddx4* expression, yielding 1,914 *Map7^+/-^* and 4,745 *Map7^-/-^* germ cells (Figure S9). Because cell recovery can vary across samples, genotype comparisons focused on integrated analyses and proportion-based shifts across transcriptionally defined states. After dataset integration and unsupervised clustering, we identified 15 clusters (0–14) (Figure S10A). To define germ cell differentiation stages, we used stage-specific marker analysis and RNA velocity profiling (Figures S10B and S10C). Metascape analysis revealed distinct biological processes enriched in each cluster, aligned with the expected progression of spermatogenesis (Figure S11). Based on inferred trajectory positions, clusters were renamed Ca–Co to reflect their differentiation order (Figures 7E and S10A). We then quantified the proportion of cells in each cluster to assess genotype-specific changes.

To determine whether cluster expansion reflected aberrant meiotic entry, we examined expression of *Stra8* and *Meiosin*, key meiotic inducers^40^. Both genes were appropriately upregulated at meiotic entry and downregulated thereafter in *Map7^-/-^* germ cells, similar to *Map7^+/-^*germ cells (Figure S12A). STRA8 protein levels were similarly unchanged between genotypes (Figure S12B), indicating that meiotic initiation and early transcriptional programs remain intact. Despite appropriate meiotic entry, cytological assessment showed delayed progression in *Map7^-/-^* testes, with significantly fewer seminiferous tubules containing round spermatids compared to *Map7^+/-^* testes (Figures S12C and S12D). Consistent with this delay, cluster-level quantification revealed a marked expansion of clusters Cg and Ci in *Map7^-/-^* testes (Figure 7F). In addition, clusters Cf–Cj exhibited transcriptional silencing of sex-chromosomal genes (Figure 7F), indicating that these populations had progressed at least to the pachytene stage. To further resolve the identity of the expanded clusters Cg and Ci, we analyzed the expression of *Prelid1* and *Ccna1*, which mark early and late pachytene stages, respectively. *Prelid1* is expressed through early pachytene, and its downregulation marks the transition to mid-pachytene^41^, whereas *Ccna1* expression increases during late pachytene and remains high into diplotene^42^. *Prelid1* expression extended through Cg, indicating that Cg represents early pachytene cells expanded in *Map7^-/-^* testes (Figure 7G). *Ccna1* was elevated in Ci of *Map7^-/-^*testes, whereas in *Map7^+/-^* testes it was restricted to Cj, identifying Ci as a late pachytene population enriched in *Map7^-/-^* mice (Figure 7G). Notably, trajectory inference suggested that, in *Map7^-/-^*testes, cells within the Ci cluster may progress toward Ck (diplotene) along an alternative route with reduced representation of the Cj state observed in *Map7^+/-^* mice (Figure 7G). Together, these findings are consistent with a model in which the postnatal Sertoli apical domain provides a cytoskeletal and junctional niche that helps pace meiotic progression, linking epithelial architecture to developmental timing in the germline.

## Discussion

We identify MAP7 as a key regulator of postnatal luminal apical domain maturation in Sertoli cells, addressing a long-standing gap in how microtubules are reorganized *in vivo* to build specialized epithelial architecture during development. Although polarized Sertoli organization is essential for compartmentalizing the seminiferous epithelium and supporting germ cell development^2^, the microtubule-based mechanisms that drive this remodeling have remained poorly defined. Using a *Map7-egfp^KI^* allele, we find that MAP7 becomes enriched on apical microtubules during postnatal maturation and supports their coordinated alignment and higher-order reorganization. Rather than acting primarily as a global microtubule-stabilizing factor, MAP7 promotes bundle-scale remodeling that accompanies apical domain assembly and redistribution of F-actin. Proteomic analyses further nominate the non-muscle myosin II heavy chains MYH9 and MYH10 as MAP7-associated factors enriched in the postnatal testis, and our imaging supports a model in which MAP7-dependent microtubule organization helps establish a spatial context for luminal NMII enrichment. Together, these findings position MAP7-dependent microtubule remodeling as a core component of a postnatal program that builds luminal epithelial architecture and supports junctional compartmentalization during the first wave of spermatogenesis.

We further found that MYH9 becomes enriched at the luminal interface where apically remodeled microtubules and F-actin converge, and this spatial organization is lost in *Map7^-/-^*Sertoli cells. In the knockout, both microtubules and F-actin show altered higher-order organization, yet they still accumulate within the inner region that would normally clear to form a lumen. Notably, MYH9 does not concentrate at these ectopic assemblies, suggesting that MAP7 helps establish an apical cytoskeletal context that supports spatially restricted NMII enrichment—without implying a strict assembly order among MAP7, actin, and NMII. Spatially confined NMII accumulation is a recurring feature of epithelial morphogenesis, including the nephric duct and ureteric bud where NMII supports constriction and adhesion^43^, and dental epithelia where apical MYH9 promotes coordinated cell movements during invagination^44^. In Sertoli cells, MAP7-dependent microtubule remodeling may similarly define an apical cytoskeletal “state” that enables NMII enrichment at the luminal interface as the apical domain matures. Although the upstream cues that position NMII remain to be identified, the selective loss of luminal MYH9 enrichment in *Map7^-/-^* testes supports the broader view that microtubule-driven organization is a key component of the cytoskeletal environment required for postnatal apical domain maturation.

Our findings also complement prior work emphasizing actin-based junctional regulation in Sertoli cells by defining microtubule remodeling as an additional architectural layer that helps impose spatial logic on junctional patterning during maturation^5,6^. In this view, microtubules act not only as structural scaffolds but also as organizing elements that may bias where junctional complexes are preferentially assembled and maintained. Because the BTB operates as an integrated junctional system rather than a single molecular entity, mismatched spatial arrangement among junctional components could be sufficient to compromise epithelial compartmentalization even when individual markers remain detectable. MAP7-dependent microtubule remodeling therefore offers a mechanistic framework linking apical cytoskeletal architecture to junctional organization during postnatal tissue remodeling.

A notable feature of the *Map7* phenotype is the separation between preserved Sertoli transcriptional identity and impaired tissue organization. Single-cell transcriptomics indicates that the Sertoli transcriptional state landscape is largely maintained despite profound disruption of apical domain maturation, consistent with MAP7 primarily regulating spatial and structural features of the niche epithelium rather than broadly reprogramming differentiation. This distinction has functional consequences: SSC maintenance and early spermatogonial differentiation appear relatively resilient to architectural defects, whereas meiotic progression is delayed in the absence of detectable defects in meiotic initiation or overt apoptosis. Together, these observations support a model in which the postnatal Sertoli apical domain functions as a structural niche component that helps pace pachytene progression, linking epithelial architecture to developmental timing in the germline.

An important question raised by this study is how MAP7’s Sertoli architectural functions intersect with potential germ cell–intrinsic requirements. MAP7 is also expressed in elongating spermatids, and *Map7* loss is associated with defects during spermiogenesis and a progressive reduction of late germ cell populations. By focusing on stages preceding the emergence of late spermatids, we minimized secondary consequences of spermatid deformation and captured early defects in postnatal apical domain maturation; nevertheless, phenotypes at later ages likely reflect combined contributions from Sertoli architectural failure and germ cell–autonomous MAP7 functions. In addition, earlier gene-trap models were maintained on mixed genetic backgrounds and did not directly establish complete loss of MAP7 protein^28^. Our independently generated knockout on a pure C57BL/6 background reproduces male infertility while providing a defined platform to evaluate how allele design and genetic background influence phenotypic presentation. Future work using tissue-restricted alleles will be essential to disentangle Sertoli-versus germ cell–autonomous contributions with higher precision.

Together, our results position MAP7-dependent microtubule remodeling as a key organizing element that builds postnatal Sertoli apical architecture and supports timely germ cell differentiation. By integrating *in vivo* genetics, multiscale imaging, proteomics, and single-cell transcriptomics, we define a MAP7-associated cytoskeletal framework linked to apical domain maturation and coordinated junctional patterning, while Sertoli transcriptional differentiation programs remain largely preserved. These findings support the idea that cytoskeletal architecture itself can act as a decisive regulatory layer for niche function. More broadly, our work highlights microtubule-driven remodeling as an autonomous architectural program in highly specialized epithelia, providing a conceptual framework that may extend to other niche-supporting tissues.

## Materials and Methods

### Animals

C57BL/6NJcl mice were purchased from CLEA Japan (Tokyo, Japan). We used *Map7^+/-^*, *Sox9-ires-gfp;Map7^+/-^*, and *Map7-egfp^KI^* mice, all maintained on a C57BL/6 background, for this study. The *Sox9-ires-gfp* mouse line was originally generated by Nel-Themaat *et al*.^45^. For scRNA-seq of total testis cells, *Map7^+/-^* or *Map7^-/-^* male mice at P23 were used. For scRNA-seq targeting Sertoli cells, *Sox9-ires-gfp* or *Sox9-ires-gfp; Map7^-/-^* male mice at P19 were used. Testes collected from mice at P10–P35 and at 8 weeks of age were used for histological and immunostaining analyses. Whenever possible, KO animals were compared with littermates or age-matched non-littermate controls from the same colony, unless otherwise specified. Mice were housed under standard conditions in a temperature-and light-controlled facility (12-hour light/dark cycle; 22 ± 1 °C). All animal procedures were approved by the Institutional Animal Care and Use Committee (approval numbers: A2019-127, A2021-018, A2023-036, and A2025-018) and conducted in accordance with institutional guidelines.

### Cell culture

Wild-type and *MAP7-EGFP^KI^*HeLa cells were cultured in Dulbecco’s Modified Eagle Medium (DMEM, 043-30085; Fujifilm Wako Pure Chemical, Osaka, Japan) supplemented with 10% fetal bovine serum (FBS, 35-015-CV; Corning, Corning, NY, USA) and penicillin–streptomycin (26253-84; Nacalai, Kyoto, Japan) at 37 °C in a humidified atmosphere containing 5% CO ^19^.

### crRNAs and genotyping primers

The sequences of crRNAs and genotyping primers used in this study are provided in Tables S1 and S2, respectively.

### Generation of *Map7^+/-^* mice and genotyping

*Map7^+/-^* mice were generated by introducing Cas9 protein (317-08441; NIPPON GENE, Toyama, Japan), tracrRNA (GE-002; FASMAC, Kanagawa, Japan), and synthetic crRNA (FASMAC) into C57BL/6N fertilized eggs via electroporation. To generate the *Map7* exon2–8 deletion (Ex2-8Δ) allele, synthetic crRNAs were designed to target the sequences GGACAAACTAGCAACCACTC(AGG) in exon 2 and CGCCTGAGGGCTCTGCACGA(AGG) in exon 8 of the *Map7* gene.

The electroporation mixture consisted of tracrRNA (10 μM), synthetic crRNA (10 μM), and Cas9 protein (0.1 μg/μL) in Opti-MEM I Reduced Serum Medium (31985062; Thermo Fisher Scientific, Waltham, MA, USA). Electroporation was performed using the Super Electroporator NEPA 21 (NEPA GENE, Chiba, Japan) on glass microslides with round wire electrodes (1.0 mm gap, 45-0104; BTX, Holliston, MA, USA). The procedure included four steps of square-wave pulses: (1) three poring pulses of 3 ms each at 30 V with 97 ms intervals; (2) three polarity-reversed poring pulses of 3 ms each at 30 V with 97 ms intervals; (3) five transfer pulses of 50 ms each at 4 V with 50 ms intervals and 40% decay in voltage per pulse; and (4) five polarity-reversed transfer pulses of 50 ms each at 4 V with 50 ms intervals and 40% decay in voltage per pulse.

The targeted *Map7* Ex2-8Δ allele in F0 mice was identified via PCR using the following primers: *Map7*_C1F (5′-TAATGCAGGACCAGTCACTCTGCTGAACAG-3′) and *Map7*_C2R (5′-CTGTCACCTGATCTTGTACCCA-3′) for the KO allele (366 bp); *Map7*_C1F and *Map7*_C1R (5′-TAAGAACAGTAACTGCCCTTGCAGAGGACC-3′) for the exon 2 of the wild-type allele (597 bp); or *Map7*_C2F (5′-ACCTGCACTCTAGTTATCCCCA-3′) and *Map7*_C2R for the exon 8 of the wild-type allele (228 bp). PCR amplicons were confirmed using Sanger sequencing. Genotyping PCR was routinely performed as follows. Genomic DNA was prepared by incubating a small piece of the cut toe in 180 µL of 50 mM NaOH at 95 °C for 15 min, followed by neutralization with 20 µL of 1 M Tris-HCl (pH 8.0). After centrifugation for 20 min, 1 µL of the resulting DNA solution was used as the PCR template. Each reaction (8 µL total volume) contained 4 µL of Quick Taq HS DyeMix (DTM-101; Toyobo, Osaka, Japan) and a primer mix. PCR cycling conditions were as follows: 94 °C for 2 min; 35 cycles of 94 °C for 30 s, 65 °C for 30 s, and 72 °C for 1 min; followed by a final extension at 72 °C for 2 min and a hold at 4 °C. PCR products were analyzed using agarose gel electrophoresis. This protocol was also applied to other mouse lines and alleles generated in this study.

### Generation of *Map7-egfp* knock-in mice and genotyping

*Map7-egfp^KI^* mice were generated using the PITCh system, as previously described^46^. Cas9 protein, tracrRNA, synthetic crRNA, and the PITCh vector were introduced into C57BL/6N fertilized eggs via electroporation. The synthetic crRNA was designed to target the sequence ACTCATATAACTTCTACATG(AGG) within the *Map7* gene and GTGCTTCGATATCGATCGTT(TGG) within the PITCh vector.

The targeted *Map7-egfp^KI^* allele in F0 mice was identified by PCR using the following primers: *Map7-egfp*_C1F (5′-CTGACCTGTTCTTCCTACAGCA-3′) and *Map7-egfp*_C1R (5′-AGCCTGAAGTTCCATTCATTGT-3′) to amplify both the *Map7-egfp^KI^* allele (996 bp) and the wild-type allele (228 bp). To confirm 5′ integration of the *Map7-egfp^KI^* allele, primers *Map7-egfp*_C1F and *Map7-egfp*_E1R (5′-TTGTACAGCTCGTCCATGCCGAG-3′) were used to amplify a 935 bp fragment; for 3′ integration, *Map7-egfp*_E1F (5′-ACCACATGAAGCAGCACGACTTC-3′) and *Map7-egfp*_C1R were used to amplify a 548 bp fragment. All PCR amplicons were confirmed by sequencing.

### Immunoblotting

For immunoblotting, testes were lysed in Laemmli sample buffer prepared in-house. After boiling, the lysates were separated by SDS–PAGE, transferred to polyvinylidene difluoride (PVDF) membranes (MilliporeSigma, Burlington, MA, USA), and immunoblotted with antibodies. Detection was performed using Amersham ECL Prime (RPN2236; Cytiva, Tokyo, Japan), and signals were visualized using the FUSION Solo imaging system (Vilber Lourmat, Marine, France). The following primary antibodies were used: mouse anti-Actin (C4, 0869100-CF; MP Biomedicals, Irvine, CA, USA), mouse anti-Clathrin heavy chain (610500; BD Biosciences, Franklin Lakes, NJ, USA), rat anti-GFP (GF090R; Nacalai, 04404-84), rabbit anti-MAP7 (SAB1408648; Sigma-Aldrich, St. Louis, MO, USA), rabbit anti-MAP7 (C2C3, GTX120907; GeneTex, Irvine, CA, USA), and mouse anti-α-tubulin (DM1A, T6199; Sigma-Aldrich). Corresponding HRP-conjugated secondary antibodies were used for detection: goat anti-mouse IgG (12-349; Sigma-Aldrich), goat anti-rabbit IgG (12-348; Sigma-Aldrich), and goat anti-rat IgG (AP136P; Sigma-Aldrich). Detailed information for all primary and secondary antibodies is provided in Table S3.

### Histological analysis

For HE staining, testes and epididymides were fixed in 10% neutral buffered formalin or in-house–prepared Bouin’s solution, followed by paraffin embedding. Sections were cut at a thickness of 6 μm and mounted on CREST-coated glass slides (SCRE-01; Matsunami Glass, Osaka, Japan). After dehydration, sections were stained with HE using standard protocols.

For immunofluorescence, testes were fixed overnight at 4 °C in 4% paraformaldehyde in PBS, then cryoprotected by immersion in 10%, 20%, and 30% sucrose solutions at 4 °C. Tissues were briefly incubated in OCT compound (4583; Sakura Finetechnical, Tokyo, Japan) at room temperature, embedded in OCT, and snap-frozen in liquid nitrogen. Cryosections (10 μm thick) were stored at −80 °C until use. Prior to staining, sections were washed once in PBS for 10 min and twice in 0.1% Triton X-100/PBS for 10 min each. Blocking was performed using 2.5% normal donkey serum (017-000-121; Jackson ImmunoResearch Laboratories, PA, USA) and 2.5% normal goat serum (005-000-121; Jackson ImmunoResearch Laboratories) in 0.1%

Triton X-100/PBS for 1 h at room temperature. Sections were incubated with primary antibodies overnight at 4 °C and subsequently with Alexa Fluor-conjugated secondary antibodies (Thermo Fisher Scientific) for 1 h at room temperature. Detailed information on all primary and secondary antibodies is provided in Table S3. For PNA lectin and phalloidin staining, FITC-conjugated *Arachis hypogaea* lectin (1:1000, L7381; Sigma-Aldrich) and Alexa Fluor 647-conjugated phalloidin (1:1000, A2287; Thermo Fisher Scientific) were used, respectively. TUNEL staining was performed using the MEBSTAIN Apoptosis TUNEL Kit Direct (8445; MBL, Tokyo, Japan) to detect apoptotic cells. Nuclei were counterstained with DAPI (D9542; Sigma-Aldrich) for 30 min at room temperature. Imaging was performed using a Leica TCS SP8 confocal microscope, and images were processed using LAS X software (version 2, Leica Microsystems, Vienna, Austria). Objectives used were HC PL APO 40×/1.30 OIL CS2 (11506428; Leica) and HC PL APO 63×/1.40 OIL CS2 (11506350; Leica), with digital zoom applied as needed for high-magnification imaging. DAPI was detected using PMT detectors, while Alexa Fluor 488, 594, and 647 signals were captured using HyD detectors. Images were acquired in sequential mode with detector settings adjusted to prevent signal bleed-through. Laser power and exposure settings were optimized for each experiment but kept constant during acquisition to ensure comparability across samples. Image brightness was adjusted linearly within a consistent dynamic range using Fiji software (v.2.16/1.54p; NIH)^47^. For Figure 7D, cell quantification was performed using Aivia software (version 14.1; Leica). Statistical analyses and graph generation were performed using Microsoft Excel (Microsoft Office LTSC Professional Plus 2021), except for ANOVA, which was conducted in R (version 4.4.0)^48^. For ANOVA, linear models were fitted using the base **stats** package (lm function), and analysis of variance was performed with the **anova** function.

### Volume electron microscopy (array tomography)

Immediately following euthanasia, testes were dissected from animals, immersed in half-strength Karnovsky fixative (2% paraformaldehyde, 2.5% glutaraldehyde in 0.1 M phosphate buffer, pH 7.3), bisected in fixative, and fixed for 48 h. The tissue was post-fixed with 1% osmium tetroxide in buffer, dehydrated in a graded ethanol series, and embedded in EPON 812 resin using TAAB Embedding Resin Kit with DMP-30 (T004; TAAB Laboratory and microscopy, Berks, UK). After trimming to a 0.5 mm square, approximately 600 serial 50 nm ultrathin sections were collected on silicon wafers. Sections were stained with uranyl acetate and modified Sato’s lead solution^49^ for 5 min each and subjected to SEM (JSM-IT800SHL; JEOL, Tokyo, Japan) to generate a stack of serial section images. Imaging was conducted at a pixel resolution of 5 nm × 5 nm, with two montaged images per section, each at 7680 × 5760 pixels. After alignment using Stacker Neo (version 3.5.3.0; JEOL), 1/2 or 1/4 binning was applied. Three-dimensional ultrastructural visualization and analysis were performed using Amira 2022 (version 2022.1; Thermo Fisher Scientific) and Microscopy Image Browser (version 2.91)^50^. For Figure 3C, regions of interest within the image stack were subsequently re-examined at a higher pixel resolution of 1.25 nm × 1.25 nm.

### Confocal imaging of MAP7-EGFP^KI^ in living seminiferous tubules

Seminiferous tubules were freshly dissected from P10 or P17 *Map7-egfp^KI^* mice and dissociated in PBS. Individual tubules were immediately transferred to 35 mm glass-bottom dishes (D11130H; Matsunami) for imaging. Focusing on the apical side allowed visualization of MAP7-EGFP^KI^ localization in its native orientation. Confocal imaging was performed using a Leica TCS SP8 microscope equipped with an HC PL APO 63×/1.40 OIL CS2 objective (11506350; Leica). Digital zoom was applied as necessary to achieve high-magnification images. GFP signals were detected using HyD detectors under standard settings.

### Immunoprecipitation

Wild-type and *MAP7-egfp^KI^* HeLa cells were lysed in a buffer containing 10 mM Tris-HCl (pH 7.5), 150 mM NaCl, 0.5 mM EDTA, and 0.5% NP-40, supplemented with complete protease inhibitor (4693116001; Roche Diagnostics, Mannheim, Germany). Testes were harvested from male C57BL/6 wild-type or *Map7-egfp^KI^* mice at P17–P20, detunicated, and homogenized in the same lysis buffer with protease inhibitors. Lysates were cleared by centrifugation at 100,000 × *g* for 10 min at 4 °C. To reduce nonspecific interactions, supernatants were precleared through incubation with 50 μL of ChromoTek control magnetic agarose beads (bmab; Proteintech, Rosemont, IL, USA) for 1 h at 4 °C with gentle rotation. The resulting supernatants were used as input material for immunoprecipitation. To isolate Map7/MAP7-EGFP^KI^ complexes, 50 μL of ChromoTek GFP-Trap magnetic agarose beads (gtma; Proteintech) were added to the lysates. After incubation, the beads were washed thoroughly with lysis buffer, and bound proteins were eluted with 40 µL of elution buffer (100 mM Glycine-HCl, pH 2.5, 150 mM NaCl) and neutralized with 4 µL of 1 M Tris-HCl (pH 8.0).

### Mass spectrometry

Immunoprecipitated proteins were separated on 4–12% NuPAGE gels (NP0321PK2; Thermo Fisher Scientific) and electrophoresed until they migrated approximately 1 cm from the loading wells. Gels were stained with SimplyBlue SafeStain (LC6060; Thermo Fisher Scientific), and protein-containing bands were excised and diced into approximately 1 mm pieces. Gel fragments were subjected to in-gel digestion: proteins were reduced with dithiothreitol (DTT; Thermo Fisher Scientific), alkylated with iodoacetamide (Thermo Fisher Scientific), and digested overnight at 37 °C using trypsin and Lysyl endopeptidase (Promega, Madison, WI, USA) in 40 mM ammonium bicarbonate buffer (pH 8.0).

The generated peptides were analyzed using an Advanced UHPLC system (ABRME1ichrom Bioscience) coupled to a Q Exactive mass spectrometer (Thermo Fisher Scientific). Raw data were acquired using Xcalibur software (version 4.0.27.19; Thermo Fisher Scientific) and processed using Proteome Discoverer version 1.4 (Thermo Fisher Scientific) in combination with Mascot version 2.5 (Matrix Science, Boston, MA, USA). Searches were conducted against the SwissProt database^51^, restricted to entries for *Homo sapiens* or *Mus musculus*. A decoy database comprising randomized or reversed sequences was used to estimate the false discovery rate (FDR), and peptide identification confidence was evaluated using the Percolator algorithm. Peptide-spectrum matches were filtered to maintain a global FDR below 1%, ensuring high-confidence identifications. The complete LC-MS/MS dataset has been deposited in jPOST^52^ under the accession numbers PXD066392 (ProteomeXchange) and JPST003958 (jPOST).

### scRNA-seq

Testes were isolated from *Sox9-ires-gfp* and *Sox9-ires-gfp;Map7^-/-^* mice at P19, or from *Map7^+/-^*and *Map7^-/-^*mice at P23. The tissue was finely minced and digested with 1.2 mg/mL type I collagenase (Fujifilm Wako Pure Chemical) at 37 °C for 10 min, followed by centrifugation to collect tissue pellets. Pellets were incubated with Accutase (Innovative Cell Technologies, San Diego, CA, USA) for 5 min at 37 °C. The digestion was terminated by adding DMEM supplemented with 10% FBS, and the resulting cell aggregates were gently dissociated by pipetting. Cell suspensions were sequentially passed through 95 μm and 35 μm strainers (BD Biosciences, Milpitas, CA, USA) to remove debris and large clumps. Cells were pelleted by centrifugation, resuspended in PBS containing 0.1% bovine serum albumin, and sorted using a SH800 cell sorter (SONY, Tokyo, Japan) to exclude dead cells. GFP-positive cells were isolated for Sertoli cell-specific scRNA-seq. Viable cells were resuspended in DMEM with 10% FBS. Approximately 7,000 single cells were loaded onto a Chromium controller (10x Genomics, Pleasanton, CA, USA) for droplet-based single-cell capture. Libraries were prepared using the Chromium Single Cell 3J Reagent Kits v3 (PN-1000075; 10x Genomics), following the manufacturer’s protocol, and sequenced on an Illumina HiSeq X platform to generate 150 bp paired-end reads. Details on the number of mice used, total cells captured, mean reads per cell, sequencing saturation, number of genes detected per cell, median UMI counts per cell, and total cell numbers before and after quality control are provided in Figures S6B and S9A. Sequencing data have been deposited in the DDBJ Sequence Read Archive under accession number DRR709131, DRR709132, DRR709133, and DRR709134.

### Statistical analysis of scRNA-seq

Raw FASTQ files were processed and mapped to the mouse reference genome (mm10, GENCODE vM23/Ensembl 98) using the Cell Ranger pipeline (version 4.0, 10x Genomics). Subsequent analyses were performed using R (version 3.6.2) within the RStudio^53^ environment (version 1.2.1335). Quality control and primary data processing were conducted using the ‘Seurat’ package (version 3.2.2) in R^54,55^. Cells expressing fewer than 200 genes were excluded to eliminate low-quality cells. Datasets were merged and normalized using Seurat’s SCTransform function. Dimensionality reduction and clustering were performed using RunUMAP, with clustering based on FindNeighbors and FindClusters functions (default parameters). Differentially expressed genes were identified using FindAllMarkers (only.pos = TRUE, min.pct = 0.25, logfc.threshold = 0.25). FindMarkers was used with default parameters for specific pairwise comparisons.

Sertoli cell populations were isolated based on expression of gonadal supporting cell markers (*Gata4, Sox9, Kitl*, and *Dhh*). Selected cells were visualized using UMAP plots, and clusters expressing markers of other gonadal lineages or high mitochondrial gene content were excluded. Remaining cells were retained for downstream analysis as Sertoli cells.

Germ cells were selected based on *Ddx4* expression and re-visualized on UMAP plots. Clusters expressing somatic markers or showing high mitochondrial gene content were excluded. The remaining cells were analyzed as germ cells. RNA velocity analysis was performed using the Velocyto package (Python 3.7.3) following default settings^56^ and visualized using UMAP embeddings generated by Seurat.

### Gene enrichment analysis

Gene enrichment analyses were performed using Metascape^57^ (v3.5.20250701) with default settings, and results were visualized in Microsoft Excel. To characterize each cluster, we used the top 100 representative genes per cluster.

### Statistics

All experiments were repeated at least three times using biological replicates. Data are presented as average ± standard deviation (S.D.) or, for box-and-whisker plots, as median, first and third quartiles, and 5th–95th percentiles. Statistical comparisons between groups were primarily performed using the Student’s *t*-test. A *P* value <0.05 was considered statistically significant. *Z*-scores for IP-MS analyses were calculated from Mascot scores. Spearman’s correlation coefficients were computed to estimate transcriptional similarity between clusters. *P* values and adjusted *P* values (*padj*) during the estimation of Sertoli cell proportions per cluster were calculated using Fisher’s exact test. One-way ANOVA was performed to assess statistical significance during the quantification of round spermatids and pachytene spermatocytes.

## Acknowledgments

We thank Dr. Nishinakamura for generously providing the MYH9 antibody. We are also grateful to members of our laboratory for their insightful discussions and technical support. This work was partially conducted at the Center for Animal Resources and Development and the Research Facilities of the School of Medicine, Kumamoto University.

## Funding

This study was supported by the Research Program for the Inter-University Research Network for High Depth Omics and the Joint Usage/Research Program for Developmental Medicine, IMEG, Kumamoto University. Additional support was provided by JSPS KAKENHI (Grant Numbers 19K06664, 22K06226, 25K09632, JP16H06280 [ABiS], JP22H04922 [ABiS], and JP16H06276 [AdAMS]); the MEXT Promotion of Development of a Joint Usage/Research System Project: Coalition of Universities for Research Excellence (CURE) Program (Grant Number JPMXP1323015486); and grants from the Narishige Zoological Science Award, the Novartis Foundation for the Promotion of Science, the Mochida Memorial Foundation, the Takeda Science Foundation, and the Uehara Memorial Foundation (to K.K.). Further support was provided by MEXT KAKENHI (JP23H00379, 25H01304), AMED PRIME (21gm6310021h0003), and grants from the Naito Foundation, the Astellas Foundation, and the Takeda Science Foundation (to K.I.).

## Author contributions

K.K. designed the study, performed the majority of the experiments, interpreted the data, and wrote the manuscript. T.F. and N.M. contributed to the initial analysis of *Map7* KO mice. K.O. assisted with array tomography analysis. R.S. contributed to scRNA-seq analysis. S.F. supported histological analysis. K.Y. and S.U. assisted with scRNA-seq analyses. N.T. conducted the mass spectrometry analysis. A.N. provided advice on IP-MS analysis. K.A. generated the knock-in mice. K.I. provided technical assistance for mouse testis analyses. All authors discussed the results and reviewed the manuscript.

## Conflict of Interest

The authors declare that the research was conducted in the absence of any commercial or financial relationships that could be construed as a potential conflict of interest.

## Data and materials availability

The LC-MS/MS and scRNA-seq datasets generated in this study have been deposited in public repositories. Accession numbers are listed in the Materials and Methods section of the main text.

## Notes

### Competing Interest Statement

The authors have declared no competing interest.

### Summary of Updates

To comply with the submission requirements, several adjustments to the manuscript were necessary. We have therefore revised and re-uploaded the manuscript text.

