## Supplementary Information for "MAP7-driven microtubule remodeling builds the Sertoli apical domain that supports timely meiotic progression"

**Figure S1. Generation of *Map7-egfp* knock-in mice.**

(**A**) Schematic representation of the generation of *Map7-egfp* knock-in (*Map7-egfp^KI^*) mice using the PITCh system.

(**B**) Genotyping of *Map7-egfp^KI^* mice. Left panel, schematic representation of the primer positions used for genotyping. Middle panel, PCR results from the three primer sets shown in the left panel. Right panel, the distinction between the wild-type (228 bp) and *Map7-egfp^KI^* (548 bp) alleles using the indicated primer sets.

(**C**) Confirmation of MAP7-EGFP^KI^ expression by immunoblotting. Lysates from the indicated mice were probed with anti-MAP7 or anti-GFP antibodies. The blot was reprobed for Actin as a loading control.

(**D**) Whole-mount immunofluorescence of fallopian tubes from *Map7-egfp^KI^* mice stained with GFP and phalloidin (F-actin). O, ovary side; U, uterus side. Asterisks indicate secretory cells that lack planar cell polarity. Scale bar, 5 μm.

**Figure S2. Generation of *Map7* knockout mice.**

(**A**) Schematic representation of the generation of *Map7* knockout (KO) mice using the CRISPR-Cas9 technique, with primer positions used for genotyping indicated.

(**B**) Confirmation of the *Map7* KO (Δ) allele by Sanger sequencing. The junction indicates the site where the two gRNA-induced cuts were ligated.

(**C**) Genotyping of *Map7* KO mice. Left panel, PCR results using three primer sets: C1F-C1R (5′: 597 bp), C2F-C2R (3′: 228 bp), or C1F-C2R (Δ: 366 bp). Right panel, discrimination between the wild-type (228 bp) and Δ (366 bp) alleles using the indicated primer sets.

(**D**) *Map7* knockout was confirmed by immunoblotting. Testis lysates from the indicated mice were probed with two anti-MAP7 antibodies, targeting either the C-terminal region (GeneTex, C2C3) or the full-length protein (Sigma-Aldrich, SAB1408648). The blot was reprobed with antibodies against Clathrin heavy chain (HC) or α-Tubulin as loading controls.

(**E**) Testis-to-body weight ratios (mg/g) of wild-type, *Map7^+/−^*, and *Map7^−/−^* mice (8-weeks old) were calculated (average ± SD). Data were collected from seven wild-type and *Map7^+/-^* mice and six *Map7^-/-^* mice. Statistical significance is indicated by *P* < 9 × 10^-7^ (Student’s *t*-test). Scale bar, 4 mm.

(**F**) Hematoxylin–eosin (HE) staining of sections from the testes or epididymis of the indicated mice (8-weeks old). Biologically independent mice (N = 3) per genotype were examined. Scale bar, 100 μm.

**Figure S3. MAP7 is required for postnatal apical–basal organization in Sertoli cells.**

**(A)** Apical–basal organization of Sertoli cells and luminal F-actin remodeling were analyzed at postnatal day (P)10 and P17. Seminiferous tubules were stained with DAPI and SOX9 to mark Sertoli nuclei, and phalloidin was used to visualize F-actin. Scale bar, 25 μm.

**(B)** Apical–basal organization of Sertoli cells was further analyzed in 8-week-old mice of the indicated genotypes. Cross-sections were stained with DAPI and SOX9. Scale bar, 50 μm.

**(C)** The percentage of SOX9^+^ Sertoli cells per cell layer at 8 weeks of age was quantified. Data were obtained from >139 SOX9^+^ Sertoli cells per genotype (>6 images, five independent experiments) and are shown as average ± S.D. Statistical significance is indicated by *P* < 2 × 10^−4^ (Student’s *t*-test).

**(D)** Testis-to-body weight ratios (mg/g) of *Map7^+/-^* and *Map7^-/-^* mice treated with or without busulfan (45 μg/g). Data are shown as average ± S.D. (n = 4 per group).

**(E)** HE staining of testis sections from mice treated with or without busulfan. Scale bar, 50 μm.

**Figure S4. MAP7 facilitates microtubule remodeling, but not microtubule stabilization, during postnatal apical domain maturation in Sertoli cells.**

**(A)** Microtubule organization and stability in Sertoli cells, F-actin organization at apical and basal ectoplasmic specializations (ES), and adherens junctions at the basal ES (β-catenin) were examined in testes from the indicated 8-week-old mice. Seminiferous tubule sections were stained for TUBB3 (βIII-tubulin), phalloidin (F-actin), β-catenin, and DAPI. Scale bar, 50 or 25 μm.

**(B)** Tight junction formation between basement membrane–attached Sertoli cells was analyzed by SEM in testes from P10 and P21 mice. Boxes indicate Sertoli–Sertoli boundaries shown at higher magnification in Figure 4B. For consistency, images in Fig. 4B were rotated to align Sertoli–Sertoli boundaries in the same orientation. Scale bar, 1 μm.

**Figure S5.** **MYH9 localization in postnatally maturing Sertoli cells.**

Testes from P17 *Map7^+/-^* and *Map7^-/-^* mice were analyzed by immunofluorescence. Seminiferous tubule cross-sections were stained for TUBB3, MYH9, phalloidin (F-actin), and DAPI. Boxed regions 1 and 2 correspond to the magnified views shown in the indicated panels. Scale bar, 50 or 25 μm.

**Figure S6. scRNA-seq of Sertoli cells from P19 control and *Map7^-/-^* testes.**

**(A)** Fluorescence-activated cell sorting (FACS) of GFP-positive cells using the *Sox9-ires-gfp* allele. Cells were gated by FSC–SSC, dead cells were excluded by propidium iodide (PI), and GFP-positive cells were collected for scRNA-seq. The control sample (“Control; *Sox9-ires-gfp*”) consisted of testes pooled from one wild-type; *Sox9-ires-gfp* mouse and two *Map7^+/-^*; *Sox9-ires-gfp* littermates. Testes from *Map7^-/-^*; *Sox9-ires-gfp* littermates were processed in parallel.

**(B)** Summary of 10x Genomics Chromium metrics for control and *Map7^-/-^* samples at P19.

**Figure S7. Metascape gene enrichment analysis for Sertoli cell scRNA-seq clusters.**

Metascape enrichment analysis was performed on the top 100 marker genes for each Sertoli cell cluster (C0–C5) identified by scRNA-seq.

**Figure S8. Delayed differentiation into round spermatids in *Map7^-/-^* testes without increased apoptosis.**

**(A)** Testis sections from 8-week-old mice stained for PNA (acrosome), SYCP3, ZFP541, and DAPI. Scale bar, 50 or 25 μm.

**(B)** TUNEL assay on testis sections from 8-week-old mice, followed by immunostaining for SYCP3 and ZFP541. Nuclei were counterstained with DAPI. Scale bar, 50 or 25 μm.

**(C)** HE staining of testes from P23 mice. The percentage of seminiferous tubules containing a lumen was quantified. Data are from >136 tubules across three independent experiments (average ± SD). Student’s *t*-test, *P* < 2 × 10^-7^. Scale bar, 500, 100, or 50 μm.

**(D)** Three-dimensional projection images showing microtubule organization in Sertoli cells at P23. Sections were stained for TUBB3 (βIII-tubulin), phalloidin (F-actin), β-catenin, and DAPI. Scale bar, 50 or 25 μm.

**Figure S9. scRNA-seq profiles of testicular cells from P23 *Map7^+/-^* and *Map7^-/-^* testes.**

**(A)** Summary of 10x Genomics Chromium metrics for scRNA-seq of dissociated testicular cells at P23. The fraction of germ cells (*Ddx4*^+^) among all captured cells is shown in the bottom row.

**(B)** UMAPs of testicular cells from P23 *Map7^+/-^* and *Map7^-/-^* testes (top). Integrated clustering analysis of combined datasets (bottom).

**(C)** Feature plots showing marker expression across clusters: *Ddx4* (germ cells), *Aard* (germ and/or Sertoli cells), *Sox9* (Sertoli cells), *Acta2* (peritubular and/or Sertoli cells), *Kitl* (somatic/endothelial), *Hsd3b1* (endothelial/Leydig/Sertoli), *Kdr* (Leydig), *C1qb* (macrophages), and *Cd40* (B lymphocytes).

**Figure S10. scRNA-seq transcriptomic profiles of testicular germ cells from P23 *Map7^+/-^* or *Map7^-/-^* testes.**

**(A)** Germ cells were extracted from the integrated dataset for clustering analysis, and clusters 0–14 were relabeled a–o according to their inferred differentiation order.

**(B)** UMAP feature plots showing expression of key stage markers: *Zbtb16* (undifferentiated spermatogonia), *Stra8* (differentiating spermatogonia), *Meiosin* (preleptotene/pre-spermatocytes), *Hormad1* (leptotene spermatocytes), *Piwil1* (early pachytene spermatocytes), *Izumo4* (late pachytene spermatocytes), and *Acrv1* (round spermatids).

**(C)** RNA velocity vectors overlaid on the UMAP.

**Figure S11. Gene enrichment analysis using Metascape for germ cell scRNA-seq.**

Gene enrichment analysis was performed using Metascape on the top 100 genes from each germ cell cluster (Ca–Co), defined by clustering of the germ cell scRNA-seq dataset..

**Figure S12.** **Meiotic entry programs are preserved, whereas progression to round spermatids is delayed in *Map7^-/-^* testes.**

**(A)** Expression profiles of *Stra8* and *Meiosin*, key inducers of meiosis, before and after meiotic entry, based on our scRNA-seq dataset.

**(B)** Immunofluorescence of P23 testis sections at the onset of meiosis. Sections were stained for STRA8, SOX9, and DAPI. Scale bar, 500 μm.

**(C)** Immunofluorescence of P23 testis sections showing seminiferous tubules containing round spermatids at the same age as the scRNA-seq analysis. Sections were stained with peanut agglutinin (PNA), SYCP3, ZFP541, and DAPI. The asterisk (*) marks a tubule containing round spermatids. Scale bar, 50 μm.

**(D)** Percentage of seminiferous tubules containing round spermatids at P23. Data are from >51 tubules per genotype across three independent experiments and are shown as average ± S.D.

**Figure S13. Full-scan images of gel electrophoresis and immunoblot analyses.**

Full-scan images of the agarose gels shown in Figures S1B and S2C are provided in the upper and lower left panels, respectively. Full-scan images of the corresponding immunoblots shown in Figures S1C and S2D are provided in the upper and lower right panels, respectively.

**Table S1. crRNAs used in this study**

| Name | Sequence: crRNA(PAM) |
| --- | --- |
| gRNA-s1 | GTGCTTCGATATCGATCGTT(TGG) |
| gRNA-MAP7KI | ACTCATATAACTTCTACATG(AGG) |
| gRNA-MAP7KO-1 | GGACAAACTAGCAACCACTC(AGG) |
| gRNA-MAP7KO-2 | CGCCTGAGGGCTCTGCACGA(AGG) |

**Table S2. Genotyping primers used in this study**

| Name | Sequence |
| --- | --- |
| MAP7EGFP^KI^-C1F (+) | CTGACCTGTTCTTCCTACAGCA |
| MAP7EGFP^KI^-E1F (+) | ACCACATGAAGCAGCACGACTTC |
| MAP7EGFP^KI^-C1R (-) | AGCCTGAAGTTCCATTCATTGT |
| MAP7EGFP^KI^-E1R (-) | TTGTACAGCTCGTCCATGCCGAG |
| MAP7KO-C1F (+) | TAATGCAGGACCAGTCACTCTGCTGAACAG |
| MAP7KO-C1R (-) | TAAGAACAGTAACTGCCCTTGCAGAGGACC |
| MAP7KO-C2F (+) | ACCTGCACTCTAGTTATCCCCA |
| MAP7KO-C2R (-) | CTGTCACCTGATCTTGTACCCA |
| Sox9KI-F (+) | CCAGATGGACCCACCAGTATCAG |
| Sox9KI-R (-) | GGGACACTCTTGAACTAGGAGTAG |
| Sox9KI-GFP (-) | CAGCTTGCCGGTGGTGCAGATG |

**Table S3. Primary and secondary antibodies used in this study**

| Company | Name, catalog number | Used for (dilutions) |
| --- | --- | --- |
| BD Biosciences | Mouse anti-Clathrin heavy chain, 610500 | IB (1:5000) |
| Cell Signaling Technology | Rabbit anti-β-Catenin (D10A8H2), #8480 | IF (1:200) |
| GeneTex | Rabbit anti-MAP7 (C2C3), GTX120907 | IB (1:3000), IF (1:200) |
| MBL | Rabbit anti-GFP, 598 | IF (1:200) |
| Millipore | Rabbit anti-SOX9, AB5535 | IF (1:200) |
| MP Biomedicals | Mouse anti-Actin (C4), 0869100-CF | IB (1:10000) |
| Nacalai | Rat anti-GFP (GF090R), 04404-84 | IB (1:3000), IF (1:200) |
| Sigma-Aldrich | Mouse anti-Acetylated tubulin (6-11B-1), T7451 | IF (1:200) |
|  | Mouse anti-α-tubulin (DM1A), T6199 | IB (1:10000) |
|  | Mouse anti-β-III-tubulin (SDL.3D10), T8660 | IF (1:200) |
|  | Rabbit anti-MAP7, SAB1408648 | IB (1:3000) |
|  | Rabbit anti-MYH9, HPA001644 | IF (1/150) |
| Thermo Fisher Scientific | Mouse anti- ZO-1 (ZO1-1A12), #33-9100 | IF (1/100) |
| Made in-house | Guinea pig anti-STRA8 | IF (1:200) |
|  | Guinea pig anti-SYCP3 | IF (1:200) |
|  | Rabbit anti-ZFP541 | IF (1:200) |
| Sigma-Aldrich | HRP-conjugated goat anti-mouse IgG, 12-349 | IB (1:10000) |
|  | HRP-conjugated goat anti-rabbit IgG, 12-348 | IB (1:10000) |
|  | HRP-conjugated goat anti-rat IgG, AP136P | IB (1:10000) |
| Thermo Fisher Scientific | Alexa Fluor594-conjugated goat anti- Guinea pig IgG, A-11076 | IF (1:1000) |
|  | Alexa Fluor488-conjugated donkey anti-Mouse IgG, A-21202 | IF (1:1000) |
|  | Alexa Fluor594-conjugated donkey anti-Mouse IgG, A-21203 | IF (1:1000) |
|  | Alexa Fluor488-conjugated donkey anti-Rabbit IgG, A-21206 | IF (1:1000) |
|  | Alexa Fluor594-conjugated donkey anti-Rabbit IgG, A-21207 | IF (1:1000) |
|  | Alexa Fluor647-conjugated donkey anti-Rabbit IgG, A-31573 | IF (1:1000) |
|  | Alexa Fluor488-conjugated donkey anti-Rat IgG, A-21208 | IF (1:1000) |

IB, Immunoblotting; IF, Immunofluorescence
