## Supplementary figures and images for "MAP7-driven microtubule remodeling builds the Sertoli apical domain that supports timely meiotic progression"

### Figure S1

Kikuchi_Figure S1

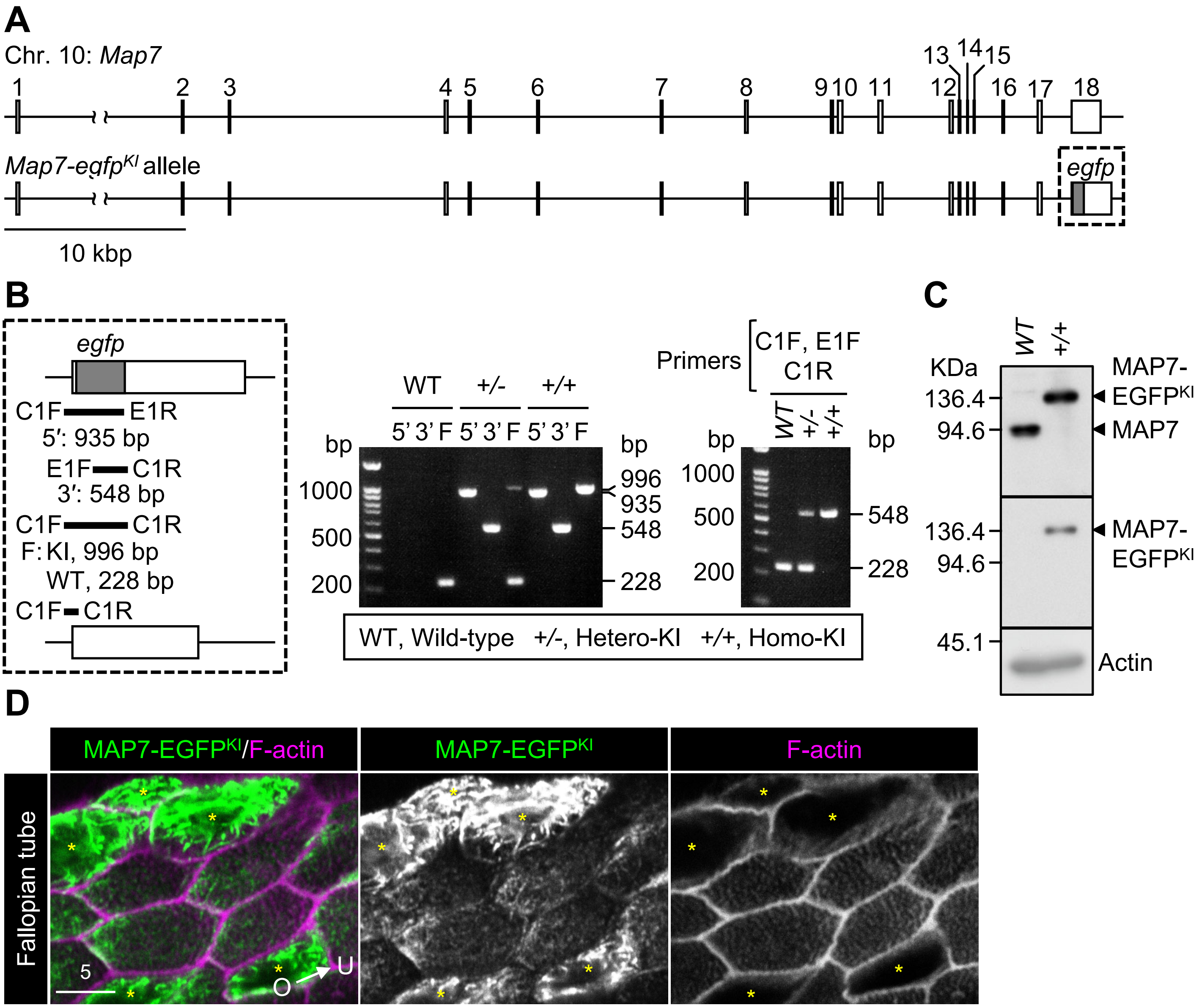

### Figure S2

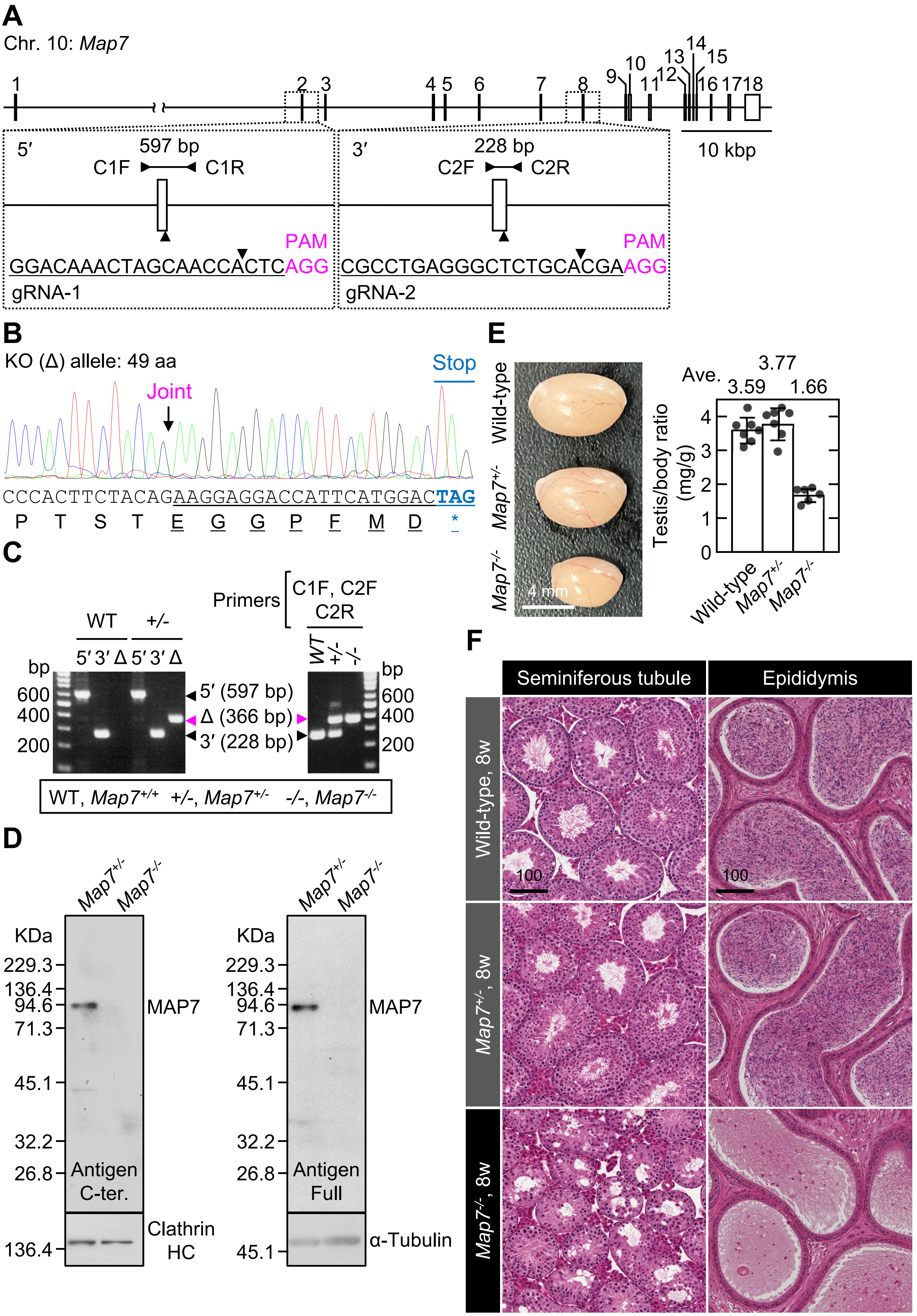

### Figure S3

Kikuchi_Figure S3

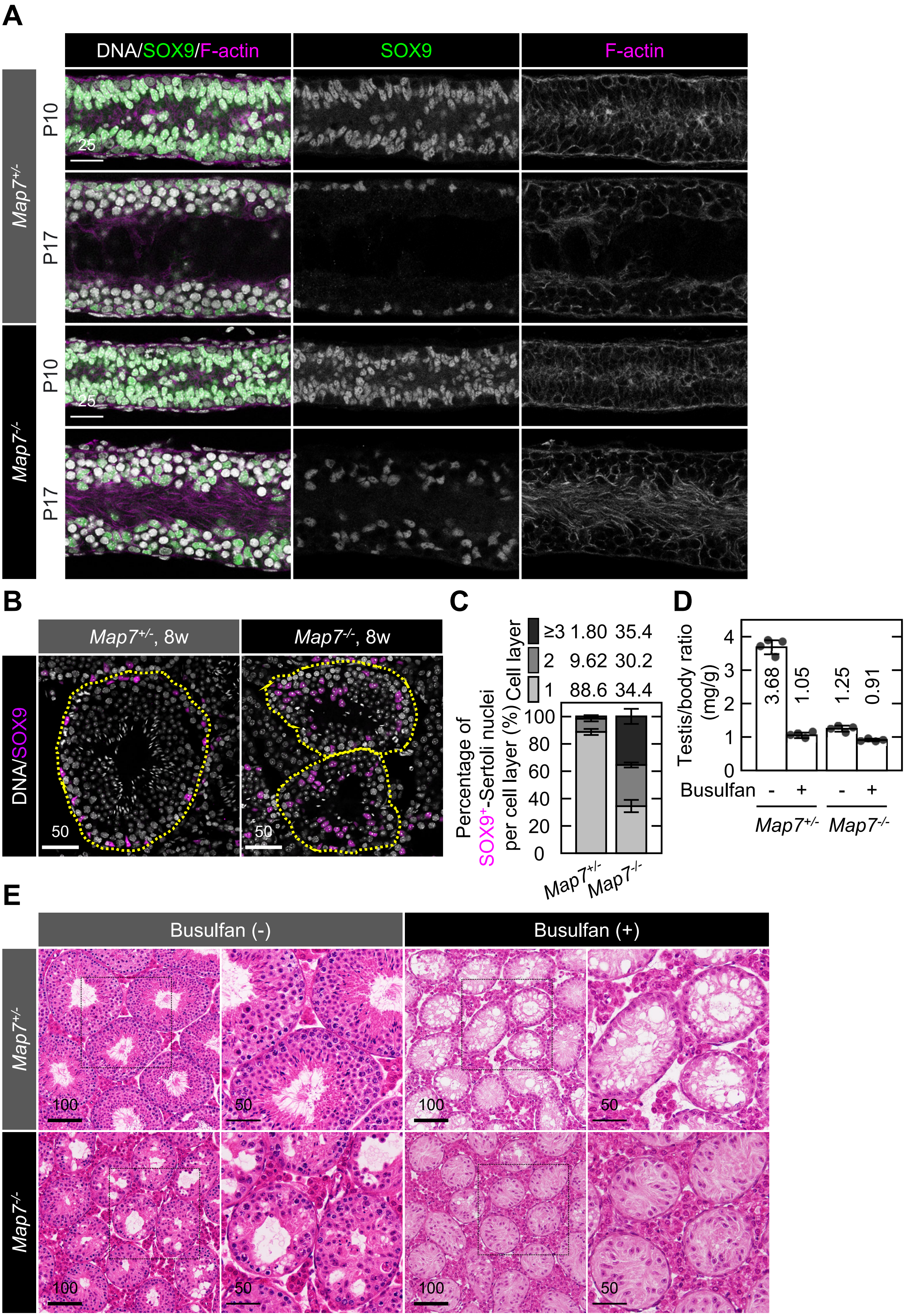

### Figure S4

Kikuchi_Figure S4

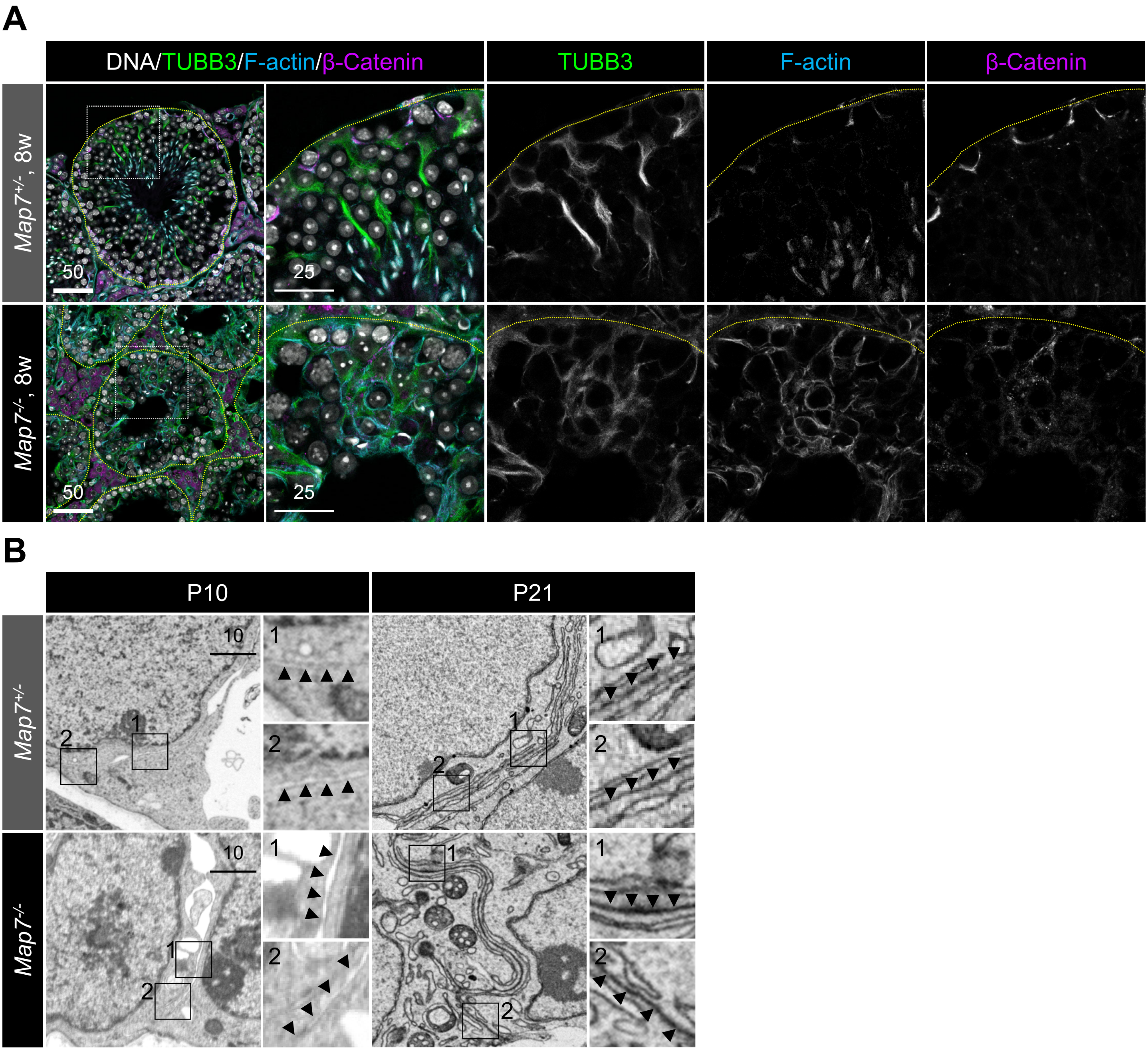

### Figure S5

Kikuchi_Figure S5

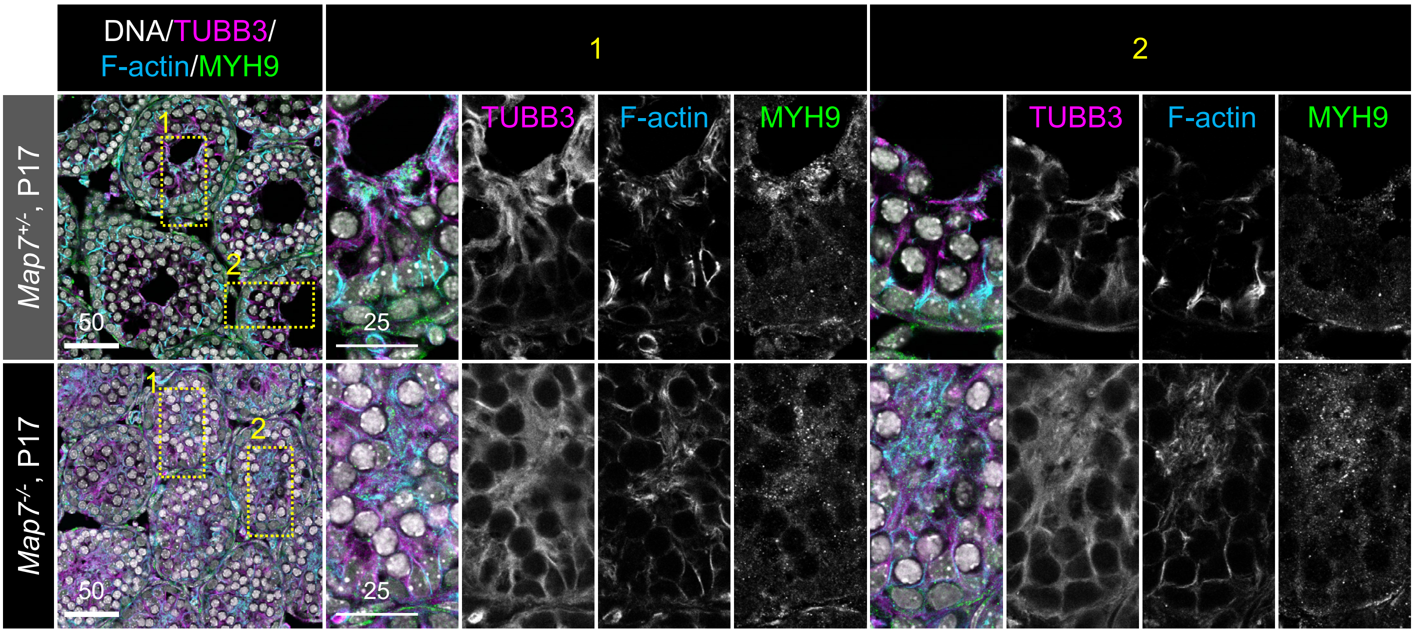

### Figure S7

Kikuchi_Figure S7

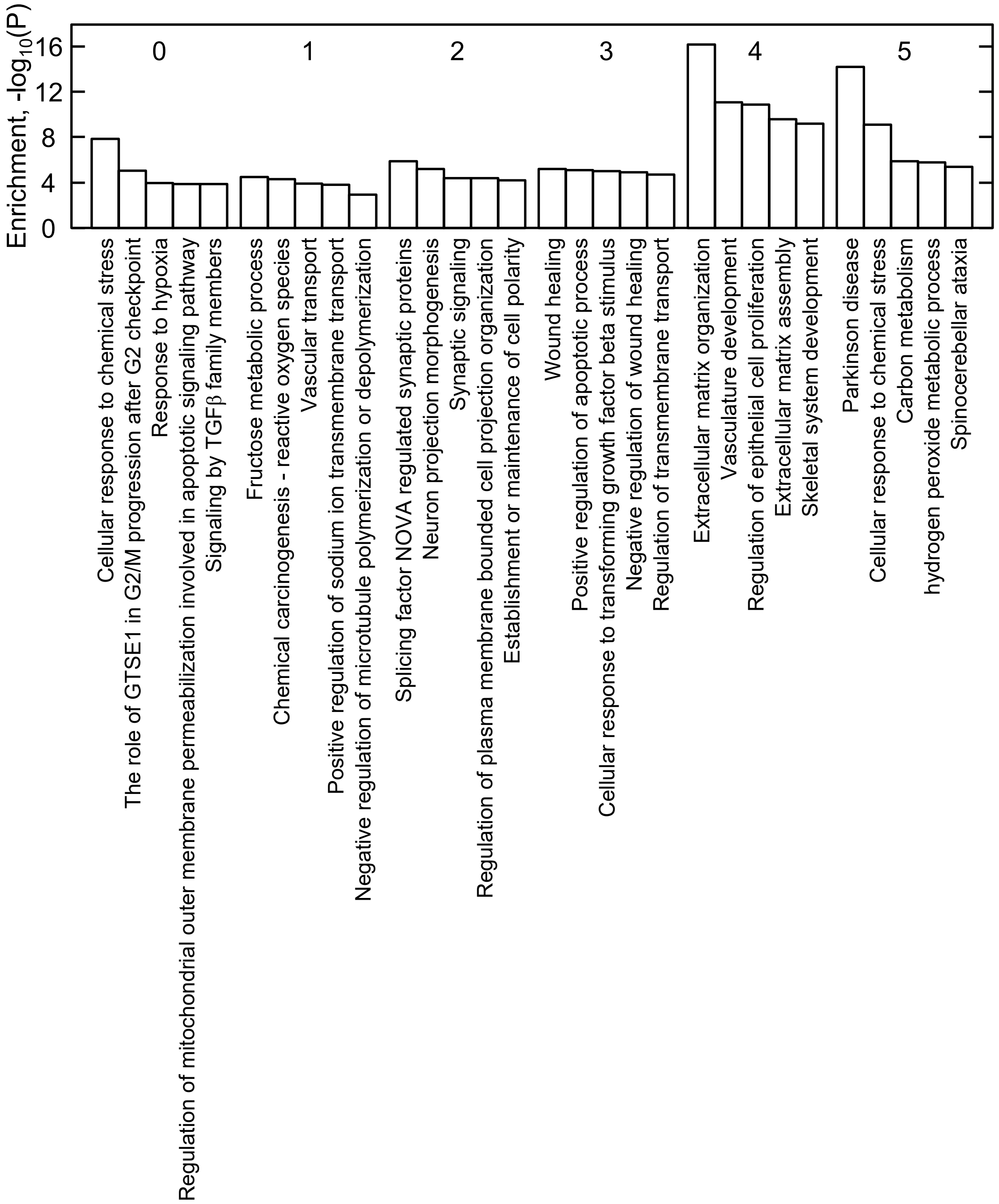

### Figure S8

Kikuchi_Figure S8

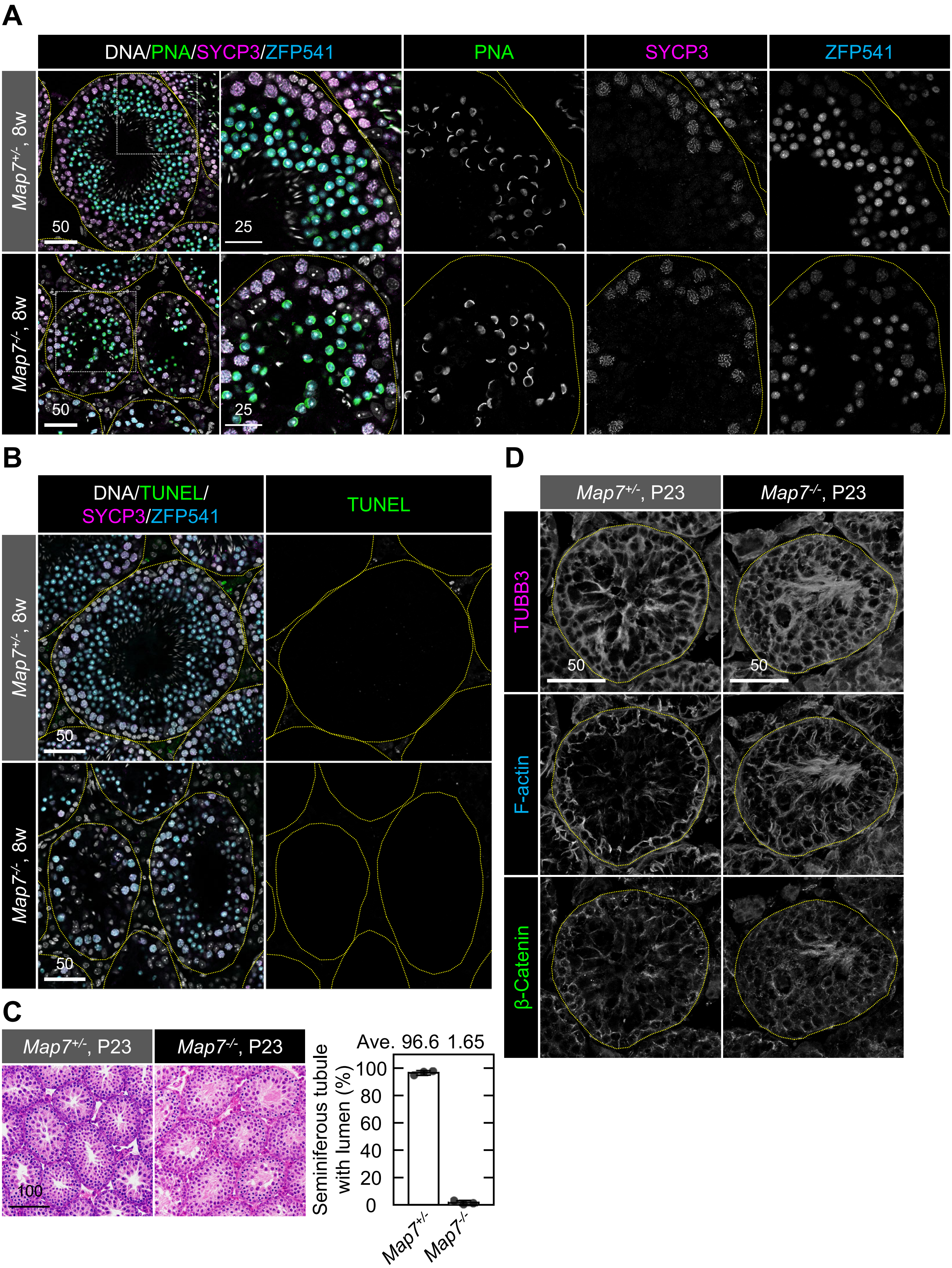

### Figure S10

Kikuchi_Figure S10

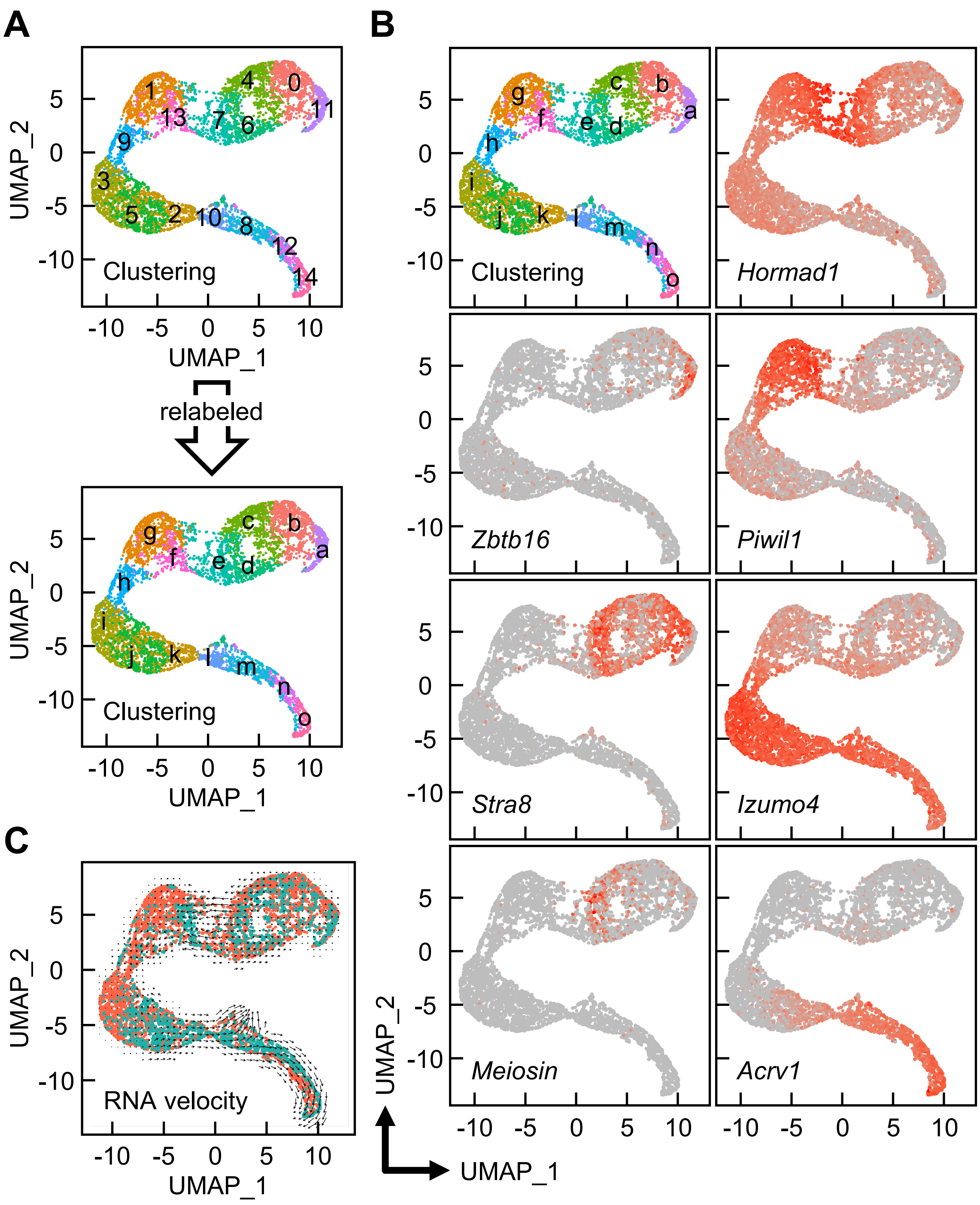

### Figure S11

Kikuchi_Figure S11

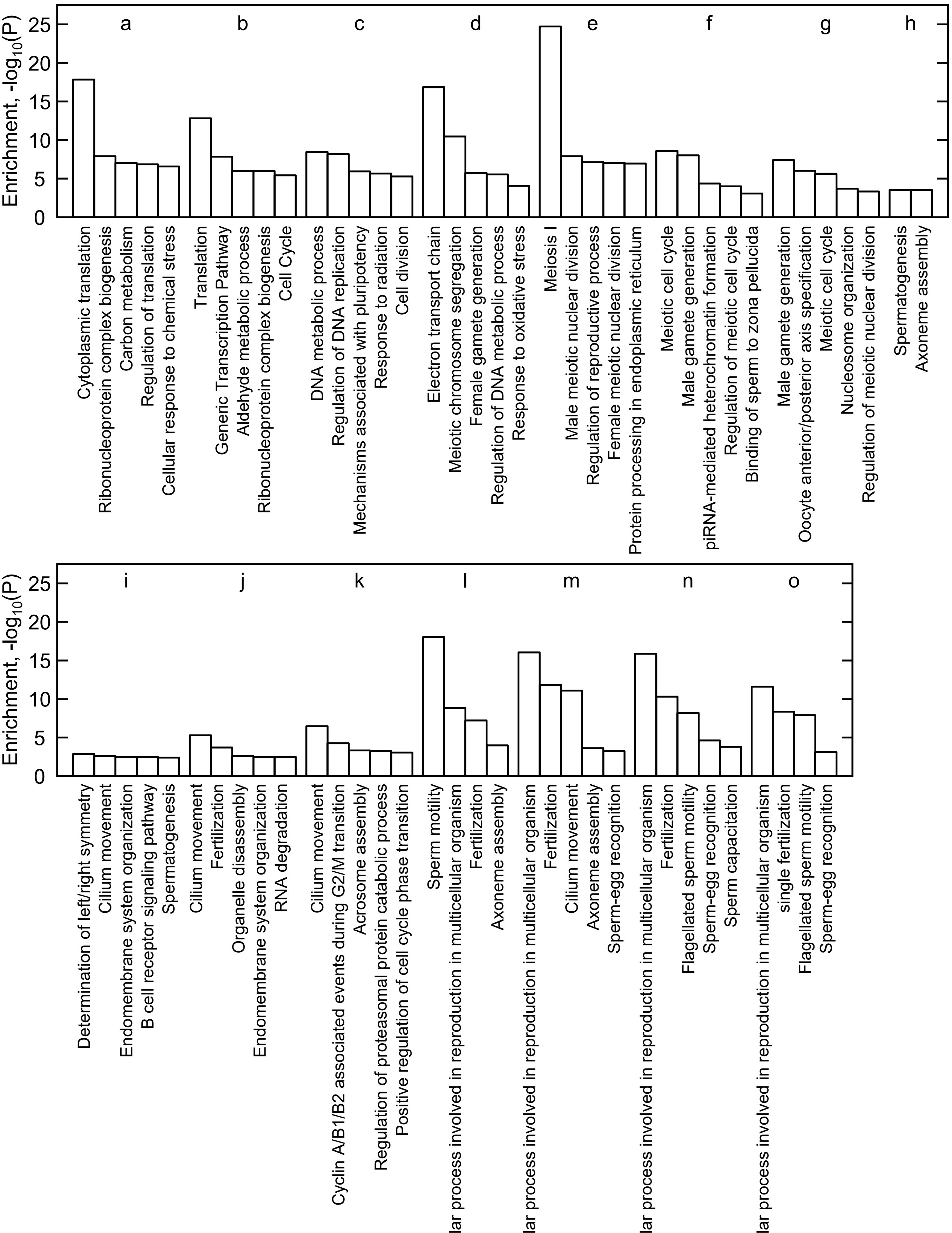

### Figure S12

Kikuchi_Figure S12

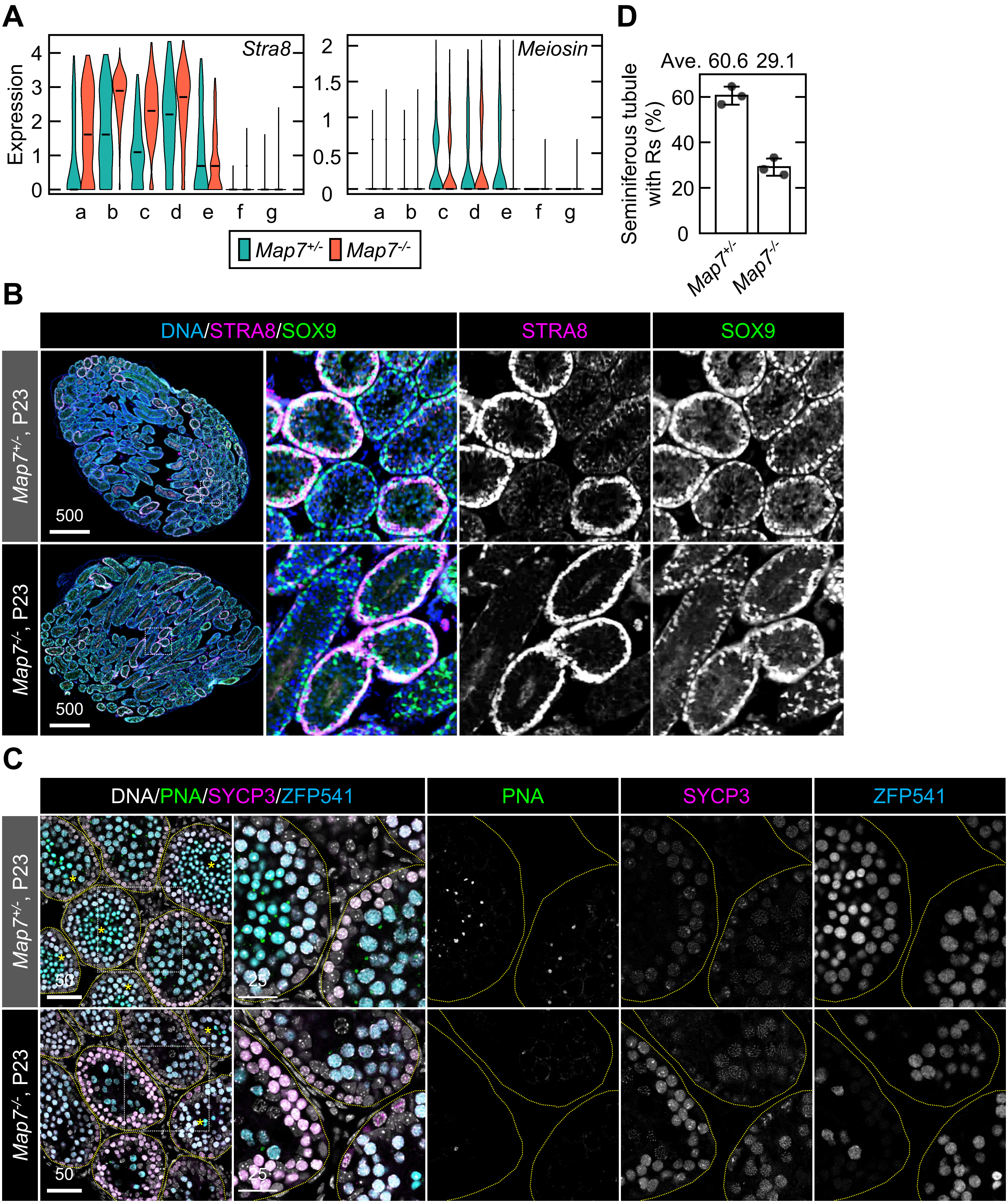

### Figure S13

Kikuchi_Figure S13

**Figure S1.**

**B**

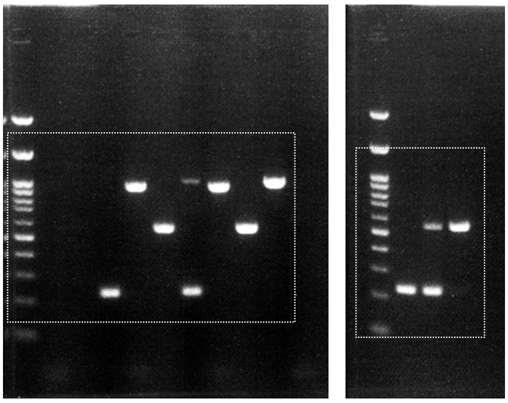

**C**

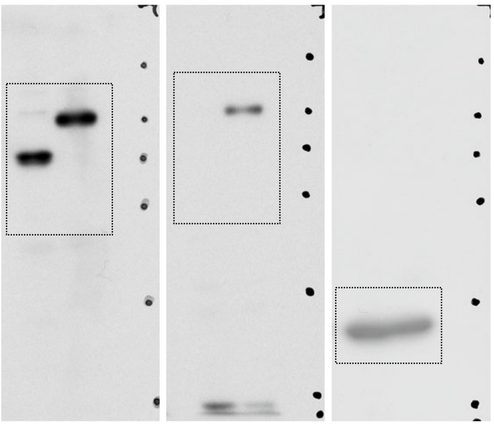

**Figure S2.**

**C**

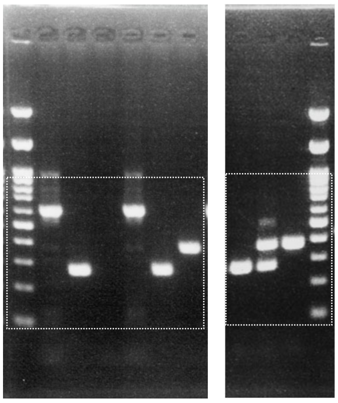

**D**

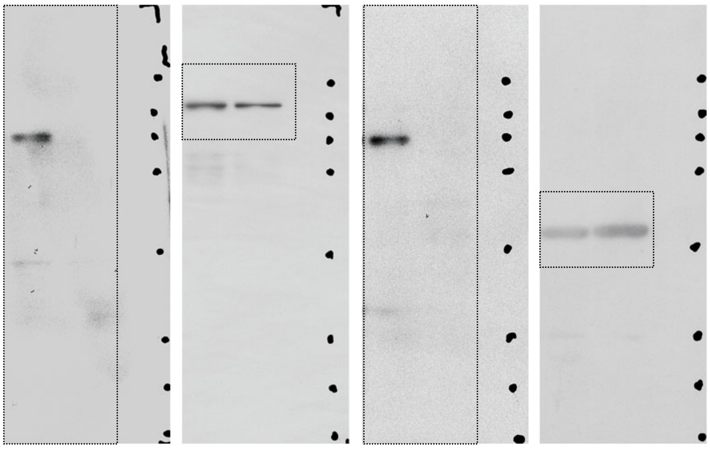
