## Supplementary material for "MAP7-driven microtubule remodeling builds the Sertoli apical domain that supports timely meiotic progression": Figure S6

Kikuchi_Figure S6

**A**

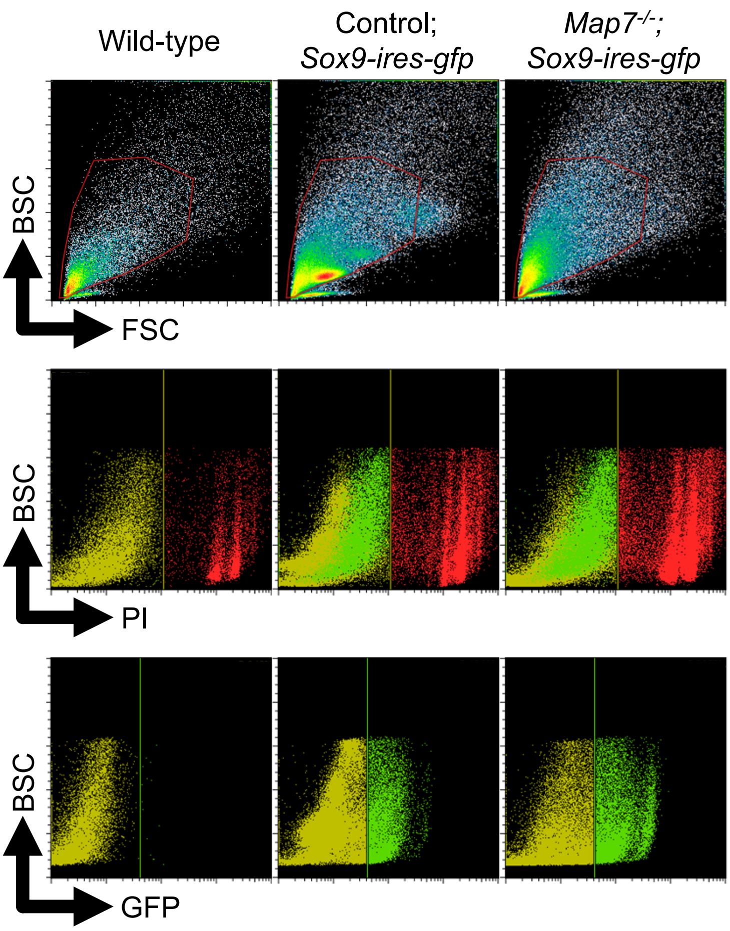


**B**

Platform: Chromium (10x Genomics)

| Sampling stage: | P19 | |
|---|---|---|
| Genotype: | Wild-type/*Map7*$^{+/-}$ | *Map7*$^{-/-}$ |
| Cell type: | GFP-sorted cells | |
| Number of used individuals: | 1/2 | 1 |
| Total number of genes detected: | 24,673 | 24,139 |
| Sequencing saturation: | 56.8% | 74.6% |
| Mean reads/cell: | 69,892 | 86,018 |
| Median genes/cell: | 2,758 | 2,256 |
| Median UMI counts/cell: | 6,471 | 5,128 |
| Total number of cells: | 5,075 | 4,535 |
| Number of analyzed Sertoli cells: | 311 | 434 |
