## Supplementary material for "MAP7-driven microtubule remodeling builds the Sertoli apical domain that supports timely meiotic progression": Figure S9

Kikuchi_Figure S9

**A**

Platform: Chromium (10x Genomics)

| | | |
|---|---|---|
| Sampling stage: | P23 | |
| Genotype: | *Map7*$^{+/-}$ | *Map7*$^{-/-}$ |
| Cell type: | Whole testicular cells | |
| Number of used individuals: | 1 | 1 |
| Total number of genes detected: | 24,847 | 25,024 |
| Sequencing saturation: | 81.0% | 89.5% |
| Mean reads/cell: | 51,385 | 34,240 |
| Median genes/cell: | 997 | 2,424 |
| Median UMI counts/cell: | 2,458 | 5,598 |
| Total number of cells: | 7,135 | 11,941 |
| Number of analyzed testicular germ cells: | 1,914 | 4,745 |

**B**

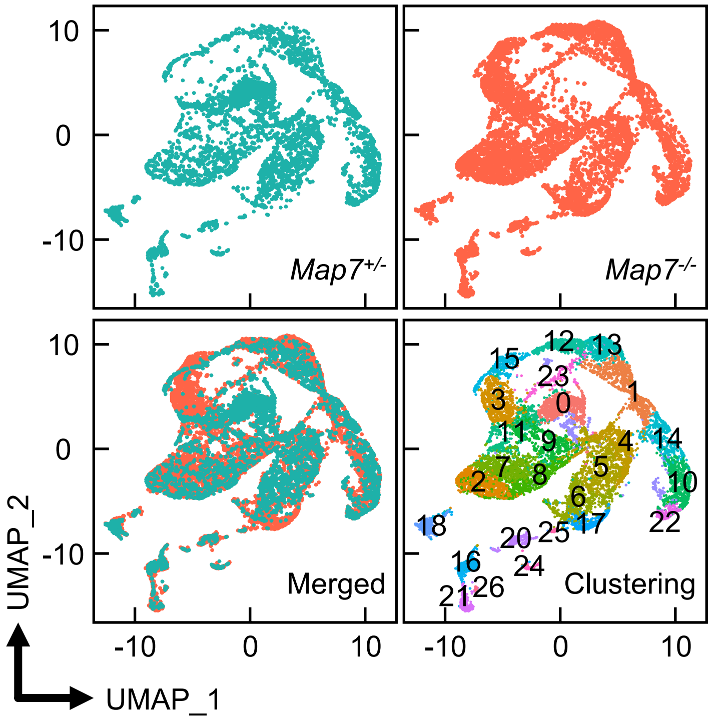


**C**

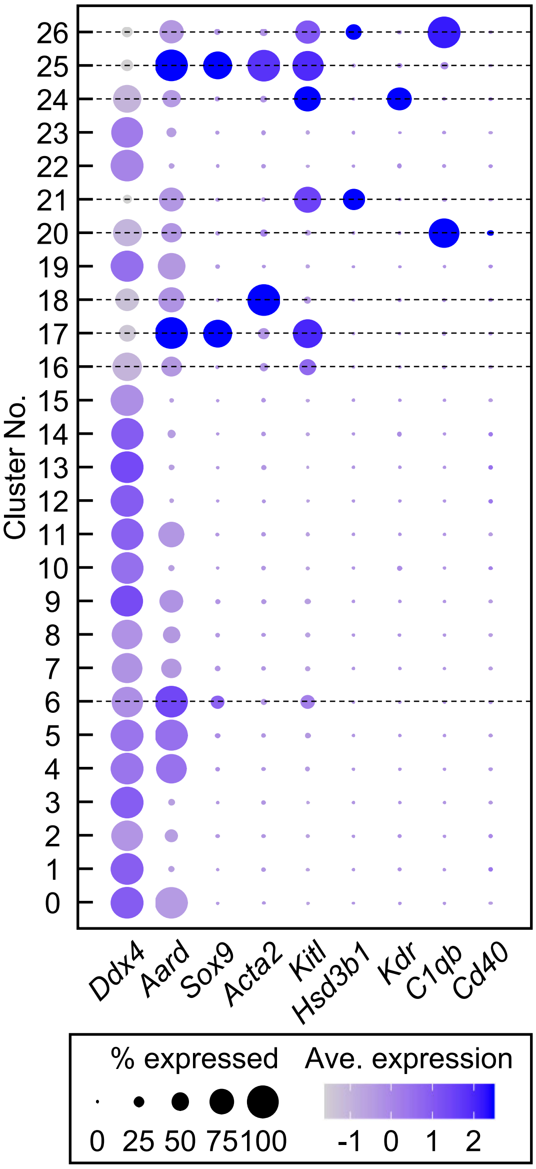
